# Antisense oligonucleotides targeting alpha-synuclein reduce pre-formed fibril-induced Lewy pathology and improve some domains of cognitive and motor performance

**DOI:** 10.1101/2020.10.21.349191

**Authors:** Sydney Weber Boutros, Jacob Raber, Vivek K. Unni

**Affiliations:** Department of Behavioral Neuroscience, OHSU; Department of Neurology, OHSU; Departments of Psychiatry and Radiation Medicine, Division of Neuroscience, ONPRC, OHSU; Jungers Center for Neuroscience Research and OHSU Parkinson Center

## Abstract

Alpha-synuclein (αsyn) is a small protein involved in neurodegenerative diseases known as synucleinopathies. The phosphorylated form (psyn) is the primary component of protein aggregates known as Lewy bodies (LBs), which are the hallmark of diseases such as Parkinson’s disease (PD) and Dementia with Lewy bodies (DLB). Synucleinopathies might spread in a prion-like fashion, leading to a progressive emergence of symptoms over time. αsyn pre-formed fibrils (PFFs) induce LB-like pathology in wild-type (WT) mice, but there are remaining questions about the progressive “spreading” of pathology and the cognitive and behavioral effects. Here, we induced LB-like pathology in the bilateral motor cortex of WT mice and assessed behavioral and cognitive performance. As there are no long-term effective treatments for synucleinopathies, and no therapies slow or reduce the spreading of LBs, we also assessed the effects of a mouse αsyn-targeted antisense oligonucleotides (ASOs) on pathology and behavioral and cognitive performance starting 5 weeks after ASO treatment. At 3 months post-PFF injection (mpi), mice injected with PFFs showed cognitive impairments and mild motor impairments. At 6 mpi, PFF-injected mice showed further cognitive and motor impairments that were partially ameliorated by the ASO. ASO treatment also reduced LB-like pathology, and pathology was significantly correlated with cognitive measures. However, the particular mouse ASO used in these assays was also associated with some possible off-target effects, defined as effects not involving lowering of αsyn, such as a decline in body weight. These results add to what is known about the progressive nature of the PFF model of synucleinopathies. These data also support the therapeutic potential of ASOs to improve Lewy pathology and associated behavioral and cognitive phenotypes.

## Introduction

Alpha synuclein (αsyn) is a small, 140-amino acid protein expressed ubiquitously in neurons throughout the central nervous system, where it is primarily localized to the presynaptic terminal and nucleus (Maroteaux, Campanelli, & Scheller, 1988; Vivacqua et al., 2011). αsyn is involved in a wide range of neural processes, such as suppressing apoptosis, regulating glucose levels, maintaining SNARE structure, and contributing to neuronal differentiation (Burre, Sharma, & Sudhof, 2014; Burre et al., 2010; Geng et al., 2011; Jin et al., 2011; Ostrerova et al., 1999; Rodriguez-Araujo et al., 2015). At the synapse, αsyn binds to vesicles and mediates membrane curvature (Wang et al., 2016). αsyn is also important for both endo-and exocytosis, and vesicle clustering; the accurate regulation of these components is essential for sustained neurotransmission, suggesting an important role for functional, monomeric αsyn (Huang et al., 2019; Jensen, Nielsen, Jakes, Dotti, & Goedert, 1998; Wu, Hamid, Shin, & Chiang, 2014; Xu et al., 2016).

Much of research into αsyn has focused on its role at the presynaptic terminal. Evidence about the nuclear function of αsyn is less clear, with conflicting conclusions about the protective or harmful nature of nuclear αsyn. αsyn associates with poly-ADP ribose, increasing pathological toxicity; inhibits histone acetylation, increasing toxicity; and protects DNA from hydroxyurea-induced stress, decreasing toxicity (Kam et al., 2018; Kontopoulos, Parvin, & Feany, 2006; Liu et al., 2011). αsyn normally binds to DNA and regulates DNA repair processes, which can be interrupted in pathological conditions (Schaser et al., 2019). However, how αsyn’s primary roles may change based on cellular and regional location are still unclear, and its wide range of functions suggest a physiological role of αsyn may be important for a healthy, functional system.

αsyn was identified as a protein of interest in neurodegenerative disease when it was discovered to be a component of senile plaques in Alzheimer’s disease, originally known as the “non-amyloid beta component,” or NAC (Ueda et al., 1993). Later, tracing of familial Parkinson’s disease (PD) led to the discovery of mutations in the gene encoding αsyn, *SNCA* (Kruger et al., 1998; Polymeropoulos et al., 1997). Subsequent investigations into PD pathology found that insoluble, phosphorylated αsyn (psyn) is the primary component of Lewy bodies (LBs) and Lewy neurites, protein aggregates that are the hallmarks of the entire class of neurodegenerative diseases now collectively known as synucleinopathies, including PD and Dementia with Lewy Bodies (DLB) (Spillantini, Crowther, Jakes, Cairns, et al., 1998; Spillantini, Crowther, Jakes, Hasegawa, & Goedert, 1998). Beyond several specific, but rare point mutations-such as A53T, A53E, A30P, E46K, H50Q and G51D-or multiplication mutations in SNCA (Appel-Cresswell et al., 2013; Fujioka et al., 2014; Kruger et al., 1998; Lemkau et al., 2012; Lesage et al., 2013; Pasanen et al., 2014; Polymeropoulos et al., 1997; Proukakis et al., 2013; Zarranz et al., 2004), the direct cause of most cases of synucleinopathies is unknown. Recent *in vitro* and *in vivo* evidence support a prion-like mechanism of αsyn aggregation, whereby introduction of exogenous αsyn pre-formed fibrils (PFFs) causes endogenous αsyn to progressively take on an insoluble, aggregate-prone conformation (Luk, Kehm, Carroll, et al., 2012; Luk, Kehm, Zhang, et al., 2012; Osterberg et al., 2015; Volpicelli-Daley et al., 2011). The idea that synucleinopathies and other neurodegenerative diseases work via a prion-like mechanism was originally put forth by Prusiner and colleagues, and has since been developed extensively (Brettschneider, Del Tredici, Lee, & Trojanowski, 2015; Brundin, Ma, & Kordower, 2016; Brundin & Melki, 2017; Prusiner, 2012; Tran et al., 2014). Importantly, monomeric forms of αsyn (Mono) do not induce LB-like aggregation, and can serve to control for the effects of αsyn over-expression (Luk, Kehm, Carroll, et al., 2012). Use of the PFF model has grown substantially as a way to study sporadic synucleinopathies since its development, but more thorough characterization of this relatively new model is needed.

Despite this progress in understanding the role of αsyn in synucleinopathies, these diseases are typically diagnosed based on symptom presentation. These symptoms often do not appear until after significant pathological spread, though. For example, PD is diagnosed based on the presence of cardinal motor signs (bradykinesia, in addition to rigidity, tremor, or postural instability) (Jankovic, 2008). Patients, however, retroactively report years of life disturbances, such as decreased olfaction (Ansari & Johnson, 1975) and sleep disturbances (Chaudhuri, Healy, Schapira, & National Institute for Clinical, 2006; Loddo et al., 2017), and continue to develop additional, non-motor symptoms, such as impaired cognition, as time passes. Post-mortem assessment of patients at various stages of disease also reveal a potential progressive spread of LBs that maps to the timing of symptom presentation (Braak et al., 2003). This likely translates to a delayed recognition of disease, to well after pathological aggregation has started and its spread has occurred.

There is currently no treatment to halt or slow the progression of synucleinopathies. Treatments are typically focused on symptom management, such as dopamine replacement therapy and deep-brain stimulation to ameliorate the motor (but not the non-motor) symptoms in PD (Del Rey et al., 2018). Additionally, treatments can have intrusive and disruptive side-effects, such as levodopa-induced dyskinesia and impulsive behavior (Pandey & Srivanitchapoom, 2017; Weintraub et al., 2006). Therefore, therapeutic strategies are being explored to decrease and/or reverse the accumulation of LBs. This has led to substantial interest in antisense oligonucleotides (ASOs), which work by binding to specific mRNAs, decreasing their translation and leading to a decrease in levels of the targeted protein. Thus, for diseases where increased levels of a known protein are thought to contribute to its pathogenicity, ASOs provide a promising avenue to reduce these levels. Currently, there are ongoing ASO clinical trials for a range of neurodegenerative diseases (Evers, Toonen, & van Roon-Mom, 2015; Schoch & Miller, 2017), including synucleinopathies (Alarcon-Aris et al., 2018; Zhao et al., 2017). Early studies have been promising, with PFF-injected mice treated with ASOs against leucine-rich repeat kinase 2 (LRRK2) showing reduced inclusion formation. However, more characterization is needed to refine treatment development.

Here, we aimed to 1) characterize the progression of behavioral and cognitive measures following bilateral injections of PFFs in the motor cortex of WT mice, and 2) assess if reducing αsyn expression throughout the CNS with a targeted ASO could ameliorate cognitive, behavioral, and pathological effects of PFF injections. We hypothesized that PFF-injected mice would show a progressive decline in cognitive and motor abilities, and that ASO treatment would reduce pathology, behavioral alterations, and cognitive impairments in PFF-injected animals.

## Materials & Methods

### Animals

These experiments involved a total of 50 mice. For the full behavioral and cognitive test battery, 35 C57BL/6J (WT) males obtained from Jackson Laboratories at 4-5 weeks of age (Bar Harbor, ME, USA) were used in all experiments. Only males were included due to the higher prevalence of PD and other Lewy-body disorders in men compared to women (Miller & Cronin-Golomb, 2010; Moisan et al., 2016). The mice were maintained on a 12hr light/dark cycle, with lights turning on at 6:00 and off at 18:00. Food and water were provided *ad libitum.*

Following three days of habituation to the new environment in the OHSU animal facility, mice underwent intracranial injection to induce LB-like pathology (“PFF”) or control (“mono”). Three months post-injection (mpi), animals underwent the first round of behavioral and cognitive testing (Fig. 1). This time was selected, as WT mice with induced LB-like pathology typically begin to show early-stage αsyn aggregation in neuritic processes at 2.5-3 mpi, as well as mild behavioral alterations, such as decreased latency to remain on the wire hang (Luk, Kehm, Carroll, et al., 2012; Osterberg et al., 2015). At 4 mpi, mice received a single, 700 μg intracerebroventricular (ICV) infusion of either an αsyn-targeted antisense oligonucleotide (“ASO”) or a scrambled control oligonucleotide (“scramble”). A second round of behavioral and cognitive testing was performed 5 weeks later to assess the effects of the targeted ASO. During periods of behavioral and cognitive testing, mice were singly housed 72 h before the start of testing; during all other periods, mice were group housed 3-5 to a cage and monitored daily for signs of fighting or distress.

**Figure 1.**
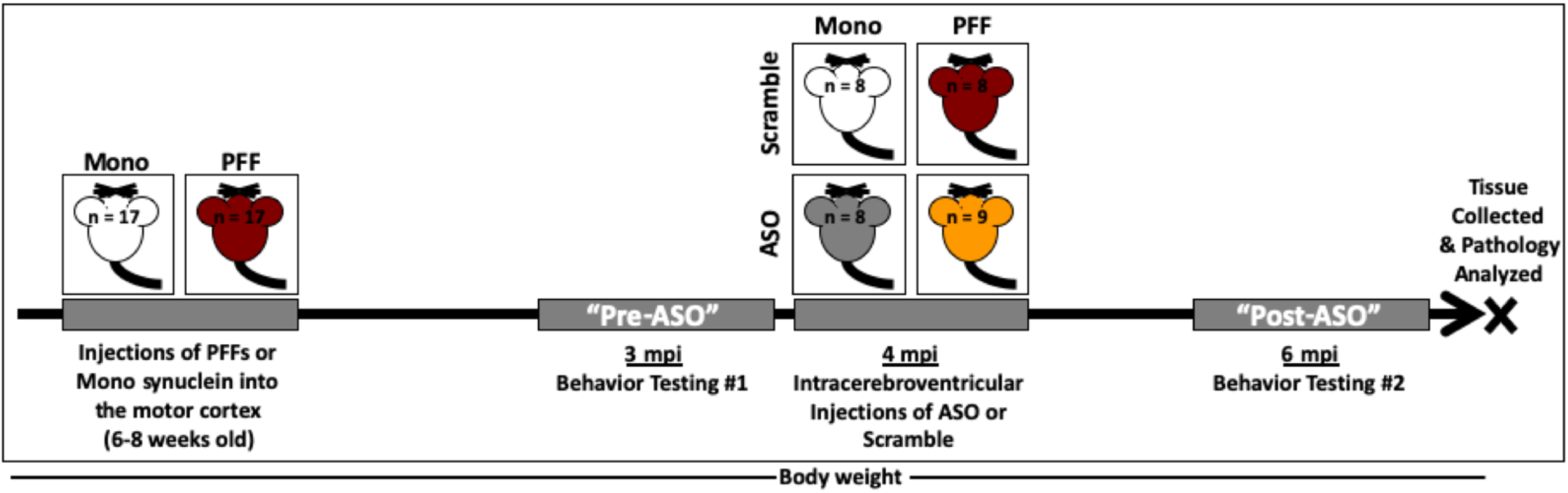
Timeline of experiments. Four-week-old C57Bl/6J male mice were delivered from Jackson Labs and acclimated to our facilities for two weeks. Mice were then injected with 2μL (2μg/μL) of either monomeric (mono) or fibrillized (PFF) a-synuclein bilaterally into the motor cortex (*n* = 17/group). At 3 months post-injection (mpi), all mice went through a battery of behavioral tests in the following order: rotarod, open field, novel object recognition, wire hang, water maze, and fear conditioning. After this round of behavior, mice received a single injection in the right ventricle with 10 μL (70 μg/μL) of ether an a-synuclein-targeting antisense oligonucleotide (ASO) or a scrambled control oligonucleotide (Scramble) (*n* = 8-9/group). Mice went through a second round of behavioral testing at 6 mpi (5 weeks following ASO delivery) in the following order: home cage activity monitoring, food intake, rotarod, open field, novel object recognition, wire hang, and water maze. Body weight was recorded weekly from the start of behavioral testing until animals were euthanized. Following completion of the second round of behavior, animals were euthanized and tissue collected for subsequent analysis.

Following the second round of behavioral testing, mice were euthanized by cervical dislocation and their brains were quickly removed. The right hemisphere was dissected into the hippocampus, cortex, and cerebellum. These tissues were flash-frozen and stored at-80° C. The left hemisphere was post-fixed overnight in 4% paraformaldehyde, and subsequently placed in 30% sucrose.

To confirm our results and assess potential off-target effects associated with the mouse ASO used in this project, an additional 15 male mice were purchased from Jackson Labs at 4-5 weeks of age. This consisted of 5 male C57BL/6N-Sncatm1Mjff/J (“αsyn-KO”), and 10 male C57Bl/6NJ (“NJ”). Half of the NJ mice received the αsyn-targeted ASO, and the other half received the control construct. All αsyn-KO mice received the ASO. None of these mice received PFF-or Mono-injections. Their body weight was tracked for 7 weeks.

All animal procedures were reviewed and approved by the Institutional Animal Care and Use Committee at the Oregon Health and Science University.

### Administered Agents & Surgeries

#### Administered Agents

##### Monomeric alpha-synuclein & pre-formed fibrils (PFFs)

Mono-and PFF synuclein were generously supplied by Dr. Kelvin Luk. Solutions for injection were prepared according to previously described protocols (Luk, Kehm, Carroll, et al., 2012). Briefly, Mono and PFFs were diluted in PBS to 2μg/μl and sonicated prior to injection as previous described (Osterberg et al., 2015). One animal was euthanized following this surgery due to recovery complications.

##### Antisense Oligonucleotides

αsyn-targeted and scrambled control antisense oligonucleotides (ASOs) were generously supplied by lonis Pharmaceuticals. αsyn-targeted ASO was matched to the non A4 component of amyloid precursor component (#678363). Mouse αsyn ASO sequence: TTTAATTACTTCCACCA; control ASO sequence: CCTATAGGACTATCCAGGAA. ASOs were diluted in 1x PBS (without Ca^2+^ or Mg^2+^) to 100 mg/ml stock solution. For injections, a working solution of 70 mg/ml was made and stored at 4°C. One animal was euthanized following this surgery due to recovery complications.

#### Surgeries

##### Motor Cortex injections of Mono/PFF

Injections of monomeric αsyn (mono) and pre-formed fibrils (PFFs) were performed according to our protocol (Osterberg et al., 2015). All mice received an injection of 0.05 μg/g buprenorphine prior to being anesthetically induced with 5% isoflurane; following induction, mice were maintained at 1.5-2% isoflurane for the duration of surgeries. Body temperature was maintained with a water heat pad, and depth of anesthesia and breathing were monitored throughout the surgeries. Following induction, mice were placed into a custom-made stereotaxic frame and heads secured with ear bars. Ophthalmic ointment was applied to the eyes to ensure they remained lubricated throughout the procedure. The heads of the mice were shaved and all mice received a local subcutaneous injection of lidocaine. The heads of the mice were sterilized by alternating 3 swabs of betadine and 2 swabs of 70% isopropanol. An incision was made on the midline of the scalp to expose the skull and holes drilled bilaterally above the motor cortex (Bregma coordinates: AP =-1.0mm, ML = ±1.5mm). A Hamilton syringe (701-RN, 26s gauge, ref# 80330) was lowered to DV-0.6mm, then raised back up to DV-0.3mm for injection of 2.5 μl (2 μg/μl) of either monomeric αsyn or PFFs at a rate of 0.2 μl/minute. The syringe remained in place for 3 minutes following infusion, and remained out for 3 minutes before lowering into the opposite hemisphere. The order of injecting into the left and right hemisphere was counterbalanced. Following injections, the scalp was stitched with two to three sutures. Animals were monitored until awake, and received 0.05μg/g buprenorphine for the following two days.

##### Intracerebroventricular infusions of ASO/Control

All ASO/Control surgeries followed the same standard procedures as above. Following placement in the stereotaxic frame, a Hamilton syringe was loaded with 10 μL of either ASO or control (70 mg/ml) and lowered into the right ventricle [AP:-0.3mm, ML: +1.0mm, DV:-2.5mm]. Infusions occurred at a rate of 2 μl/min, and the syringe was left in place for 1-minute following completion of infusion. Again, animals received 2-3 sutures and buprenorphine for post-operative care.

### Behavioral and Cognitive Assessments

#### Health Measures: Body Weight, Food Intake, & Circadian Activity

Body weights were recorded weekly starting during the first round of behavioral testing until animals were euthanized.

During the second round of behavioral and cognitive testing (6 mpi), food intake was recorded. The food in each cage was weighed twice each day-once in the morning (∼8:00) and once in the evening (∼17:00)-to assess approximate amount of food eaten during daytime hours and nighttime hours.

Additionally, home-cage activity was assessed over a week-long period as reported following ASO delivery (Torres et al., 2018). Mice were singly housed in cages containing infrared sensors, and data were continuously collected from 12:00 PM on a Monday until 12:30 PM on a Friday (MLog, Biobserve, Bonn, Germany). Following collection in 1-second increments, data were compiled into 5-min, 30-min and 12-hr bins for analyses of light and dark activity.

#### Open Field

To assess locomotion, spatial learning, and anxiety-like behaviors, mice were placed into an open field (41 × 41 cm) and allowed to explore for five minutes over three (3 mpi) or two (6 mpi) subsequent days, similar to standard protocol (McGinnis et al., 2017). The enclosures were thoroughly cleaned with 0.5% acetic acid and dried between each trial. Light intensity in the enclosures ranged from 300 to 500 lux. Animals were video recorded at a rate of 15 samples per second, and total distance moved, average velocity, and time spent in the center (defined as a center square sized 20 × 20cm) was analyzed using Ethovision XT 7.1 software (Noldus Information Technology, Wageningen, Netherlands). Data were analyzed using repeated measures ANOVAs, with PFF status and ASO treatment used as between group variables.

#### Novel Object Recognition

Following testing in the open field, two similar objects were secured in place in the center of the same fields (McGinnis et al., 2017). Animals were allowed to explore for 15 minutes (novel object day 1). The next day, one object was replaced with a distinct, novel object. Again, animals were allowed to explore for 15 minutes (novel object day 2). Total time exploring the objects was manually recorded for both sessions, as well as time spent with each individual object. The researcher recording exploration time was blinded to the groups. Percent time exploring the novel object was calculated on day 2 as an indicator that animals could distinguish which object they had previously seen. This test was repeated at both 3 mpi and 6 mpi; all the objects used at 6 mpi were distinct from the objects used during the first round of testing. Data were analyzed using repeated measures ANOVAs and two-way ANOVAs followed by Sidak’s post-hoc comparisons when appropriate.

#### Rotarod

Motor function and endurance were tested using a rotarod (Rotamex, Columbia Instruments, Columbus, Ohio) as previously reported (McGinnis et al., 2017). Mice were placed onto a rotating rod starting at 5 rotations per minute (rpm). Every three minutes, the speed increased by 1rpm. The maximum run time was capped at 300 seconds, or 50rpm. During the first round of behavioral testing, mice were tested over three consecutive days; on each day, mice received three trials, separated by three minutes each. During the second round of behavioral testing, the mice received a single day of rotarod with three trials separated by three minutes each. The rotarod was cleaned with 0.5% acetic acid between every trial. Latency to fall (in seconds) and final rpm were recorded.

#### Wire Hang

Motor function was also assessed using the wire hang task, adopting the “falls and reaches” method described by van Putten 2016 (van Putten, 2016). Mice were placed on a suspended metal wire so that they were hanging only by their front paws. In this method, mice start with a “fall score” of 10 and a “reach score” of 0. Over the duration of 180s, mice lost 1 point from the score every time they fell and gained 1 point every time they reached one of the poles holding up the wire. The time of each fall or reach event was also recorded. Each time a mouse fell or reached, the timer was paused to replace the mouse in the center of the wire again. This test has the benefit of not only allowing assessments of endurance and strength, but also of exploring more complex motor coordination.

#### Water Maze

Spatial learning and memory were assessed using the Morris water maze similar to previously described protocols (Johnson et al., 2014). A circular pool (140 cm in diameter) was filled with room temperature water (23°C ± 1) and made opaque with white chalk. The pool was divided into four quadrants (NE, NW, SE, and SW). A circular platform (12cm in diameter) was placed 2cm below the surface of the water; in order to complete the maze, mice were required to locate the platform and remain on it for a minimum of three seconds. Task learning was assessed during “visible” platform trials; a 50mL conical tube weighed down and wrapped in colorful tape was placed as a flag on top of the platform to serve as a visual cue. Spatial learning (acquisition) was assessed during “hidden” platform trials; the conical tube was removed and spatial cues of different shapes and colors were hung on the walls surrounding the pool. Training consisted of two trials per session (separated by 10 minutes) and two sessions per day (separated by 3 hours). Mice were taken out of the pool once they located and remained on the platform for 3 seconds, or after 60 seconds had passed without finding the platform. For training purposes, when mice did not find the platform on their own, the researcher guided the animals to the platform and stepped out of view for the animal to associate the platform with escape. Average velocity, cumulative distance to the target platform, and latency to find the platform were assessed as indications of learning.

To assess spatial memory retention, mice were tested in a series of probe trials in which the platform was removed from the pool. Probe trials lasted for a total of 60 seconds, and ended with the researcher picking up the animals from the location where the platform would have been located. The same extra-maze cues used during hidden trials remained on the walls for probes. Cumulative distance from the target, and time spent in the target quadrant as compared to the three non-target quadrants were analyzed as measures of memory retention.

Water maze was performed at both 3 and 6 mpi. At both time points, mice were first trained to locate a “hidden” platform. During the first round of testing, the platform remained in the SW quadrant for sessions 1-10. We adopted a reversal paradigm to challenge the mice, so for sessions 11-14, we moved the platform to a new (the opposite) quadrant (NE). Lastly, we performed a single visible platform session to ensure all mice had learned the task. During the second round, the platform was placed in the SE quadrant for sessions 1-8. On the last day (sessions 9-10), we administered two visible platform sessions with the platform in the NW and SW quadrants, respectively.

#### Fear Conditioning

Mice were tested for contextual and cued fear learning and memory at 3 mpi as previously described (Olsen, Johnson, Zuloaga, Limoli, & Raber, 2013). Briefly, mice were placed into a sound-attenuating chamber. After a 120s baseline period without any stimuli, a tone (80 dB, 2800 Hz) was played for 30s, which co-terminated with a 2s shock (0.5mA). This was followed by a 90s period with no stimuli before the tone-shock pairing was repeated again. In total, mice received 2 tone-shock pairings. The chambers were thoroughly cleaned with 0.5% acetic acid between animals. Twenty-four hours later, mice were tested for contextual and cued fear memory recall. For contextual memory recall, mice were placed into the same chamber for a period of 5 min, with no cues presented. For cued memory recall, the chambers were changed to remove any contextual cues (floors were covered with a solid, white panel, the roof and walls were changed to a black triangle, and 10% isopropanol was used for cleaning). Animals had a baseline period of 90s with no stimuli; at 90s, the same tone played for a total of 3min.

All trials were recorded and analyzed with Video Freeze software (PMED-VFC-NIR-M, Med Associates, Inc., Fairfax, VT) and automatic outputs were generated to analyze average motion and time freezing (defined as a minimum of 30 frames, or 1s, of total movement cessation). Percent time freezing was analyzed as measure of fear memory and assessed with a 1-way ANOVA between PFF and Mono groups, or with a 1-way repeated measures ANOVA when looking over time within a trial. Because we did not observe any significant differences between PFF and Mono groups at 3 mpi, we did not repeat fear conditioning testing at 6 mpi.

### Post mortem analysis

#### Immunofluorescence and microscopy

Lewy body-like aggregates were analyzed by immunofluorescence. Following post-fixation, the left hemisphere was sectioned using a cryostat at 40 μm thickness. The sections were washed, blocked in 4% normal goat serum with 0.3% Triton-X (NGS), and incubated overnight in 4% NGS containing an antibody targeting αsyn phosphorylated at serine 129 (anti-pSer129, rabbit monoclonal, 1:600, Abcam #51253). Subsequently, the sections were incubated with secondary antibody in 4% NGS overnight (goat anti-rabbit IgG, 1:1000, AlexaFluor 594, Lifetechnologies, #A11012). The nuclear counterstain DAPI (Sigma Aldrich, St. Louis, MO, USA) was applied at a 1:200 concentration for 20 minutes prior to slide mounting. Coverslips were sealed with VectaShield Hardset Antifade Mounting Media without DAPI (Vector Labs, Burlingame, CA, USA, #H-1400). Whole-hemisphere images were taken using a Zeiss AxioScan.Z1 at 20x magnification (Zeiss, Thornwood, NY, USA). The percent area occupied by immunoreactivity in the motor cortex, medial and lateral somatosensory cortices, and the hippocampus were analyzed using ImageJ Software (NIH).

#### Western blots

Western blot analysis was used to assess phosphorylated and total protein levels. Dissected hippocampi were dissolved in lysis buffer (1M Tris-Cl, pH 7.5; 6M NaCl; 10% SDS; 0.5M EDTA; Triton-X 100; Phosphatase Inhibitor #3, Roche, #05-892-970-001; Protease Inhibitor, Sigma-Aldrich, #P0044) by homogenizing 3 x 20s and sonicating on ice for 15s at 40Hz. A BCA kit (Pierce, Thermo Scientific, #23227) was used to determine protein concentrations. Samples were incubated in Novex Tris-glycine SDS Sample Buffer (Invitrogen) at 95° C for 10 min and separated on 10-20% Tris-glycine gels (Invitrogen) for 70 min at 125 V. Proteins were transferred onto Immobilon-FL PVDF membranes (Millipore, #IPF00010) for 2 h at 30 V on ice. When assessing αsyn and psyn protein levels, membranes were immediately fixed in 4% PFA + 0.1% GA for 20-30 min (Lee & Kamitani, 2011; Sasaki, Arawaka, Sato, & Kato, 2015). Total protein measures were assessed using REVERT™ Total Protein Stain (Li-Cor Biosciences, Lincoln, NE). Following image acquisition of total protein, blots were blocked in Odyssey Blocking Buffer (Li-Cor Biosciences) for 1 hr at room temperature and incubated overnight at 4° C with primary antibodies against Syn1 (mouse, 1:1000, BD Biosciences, #610786), pSer129 (rabbit, 1:1000, Abcam, #51253), GAPDH (mouse, 1:10000, Millipore, # MAB3740), or phospho-histone H2Ax (rabbit, 1:1000, Cell Signal, # 9718). Blots were then incubated in appropriate secondary antibodies (goat anti-rabbit IR800 CW, Li-Cor, #926-32211; goat anti-mouse IR680 LT, Li-Cor, #926-68050) for 2 h at room temperature. Images of hybridized blots were taken using a Li-Cor Odyssey CLx and densomitry analyses were performed using ImageJ software. The levels of the proteins of interest were normalized to the total protein loaded for that particular lane for statistical analyses.

### Statistics

All data were first assessed for normality of variance. In all cases, we were able to proceed with standard parametric tests. Statistical analyses were performed using SPSS v.25 (IBM, Armonk, NY) and Prism v.7 (GraphPad, San Diego, CA).

Upon completion of behavioral and cognitive testing, all behavioral and cognitive measures were analyzed with PFF-status (Mono/PFF) and ASO-treatment (Control/ASO) as between group variables using ANOVAs. Most measures were analyzed with a repeated measures ANOVA (body weight; home cage activity monitoring; food intake; open field activity, velocity, and center duration; time exploring during object recognition; rotarod latency to fall; wire hang latency to fall and reach, and fall and reach scores; water maze average velocity, latency to find the platform, and cumulative distance from the platform; and time freezing during each fear conditioning trial). For some tests, averages or “final scores” *(e.g.,* final fall score in wire hang) were compared with 1-or 2-way ANOVAs. For all pre-ASO tests, ASO status was dropped from the statistical model when no significance or interaction was found, and independent samples t-tests were used to compare Mono-and PFF-injected groups.

Analysis of LB-like pathology was performed with independent samples t-tests comparing ASO and control in only PFF-injected animals for each brain region. To assess overall pathology load, we normalized psyn signal in each brain region to the PFF-Scramble average of that region. We then combined regions for a composite burden measure to analyze between groups. We did this both for all regions measured, as well as with only cortical regions. Comparisons were made with independent samples t-tests between PFF-Scramble and PFF-ASO. Mono-injected animals were not included in statistical analysis, as there was no area occupied in any brain region measured for either ASO group. For Western blot analysis, bands of interest were normalized to total protein, and 2-way ANOVAs were used to compare groups.

In all cases, when significance was found with ANOVAs, post-hoc comparisons were made and Bonferonni corrections applied. In analysis of time exploring the novel vs. the familiar object, Sidak’s post-hoc comparison was applied within each group.

## Results

### PFF injections do not alter fear learning and memory

Prior to delivery of αsyn ASO or control, non-targeting ASO, we assessed all animals in a battery of behavioral and cognitive tests to determine the effects of PFF injections in the motor cortex on motor abilities, anxiety-like behavior, and learning and memory. This first round of behavioral testing took place 3 mpi, about when we expect mature LB-like inclusions to have appeared in the PFF group (Osterberg et al., 2015). Following the first round of behavioral tests, the Mono-and PFF-injected groups were split, with half in each group receiving control or targeted ASO injections (n = 8-9/group). Seven weeks after ASO delivery, animals were tested in a second battery of behavioral tests.

In the first round of testing (pre-ASO), we assessed fear learning and memory to examine possible hippocampus-dependent and-independent effects. Fear conditioning can distinguish between hippocampus-dependent and-independent learning and memory. Contextual fear conditioning is known to be hippocampus-dependent, whereas cued fear conditioning is known to be amygdala-dependent (Bocchio, Nabavi, & Capogna, 2017; Curzon, Rustay, & Browman, 2009). Pre-ASO delivery, we trained mice in a cued fear conditioning paradigm, and assessed their contextual and cued recall 24 h later. However, we did not see differences between groups in acquisition of fear, or in contextual or cued recall, indicating no fear memory impairments (Table 1). As such, we did not repeat fear conditioning in the second round of behavioral testing.

**Table 1.** Performance in the fear conditioning test at 3 mpi. No differences were detected between Mono-and PFF-injected animals in any measure. All data are presented as averages ±SEM.

| Trial | Measure | Mono | PFF | Statistical Test | *p* value |
| --- | --- | --- | --- | --- | --- |
| Training | Baseline Average Motion (au) | 245.77 | 229.94 | t-test | 0.428 |
| Training | Average Motion during Shocks (au) | 893.27 | 919.95 | RM-ANOVA | 0.708 |
| Training | % Time Freezing during Tones | 5.22 | 7.52 | RM-ANOVA | 0.403 |
| Training | % Time Freezing during ISIs | 10.21 | 11.77 | RM-ANOVA | 0.616 |
| Context | % Time Freezing Total | 18.52 | 19.47 | t-test | 0.772 |
| Cued | Baseline Average Motion (au) | 278.58 | 264.45 | t-test | 0.409 |
| Cued | % Time Freezing during Tone Total | 18.22 | 23.78 | RM-ANOVA | 0.188 |

### PFF injections are associated with hippocampus-dependent spatial learning and memory impairments at 3 mpi

During the first round of behavioral testing, we assessed hippocampus-dependent spatial learning and memory using the Morris Water Maze (WM). For the first 5 days, the platform was located in the SW quadrant with extra-maze cues hung on the wall (“hidden” platform trials); for the subsequent 2 days, the platform was moved to the NE quadrant; and for the final day, we assessed visible platform learning (Fig. 2A). We saw a pronounced deficit in PFF animals’ ability to learn to locate a hidden platform, shown by a significantly higher cumulative distance to the target platform over the first 5 days of training (repeated measures ANOVA: time x PFF interaction, *F*(4.647,144.055) = 2.322, *p* = 0.050, Greenhouse-Geisser corrected; main effect of PFF, *F*(1,31)= 9.373, *p* = 0.005; Fig. 2B). There were no differences found in average swim speeds (*p* > 0.05, all platform locations), indicating that this deficit was not due to impaired locomotion. When the hidden platform location was changed, this difference was no longer detected (Fig. 2B).

**Figure 2.**
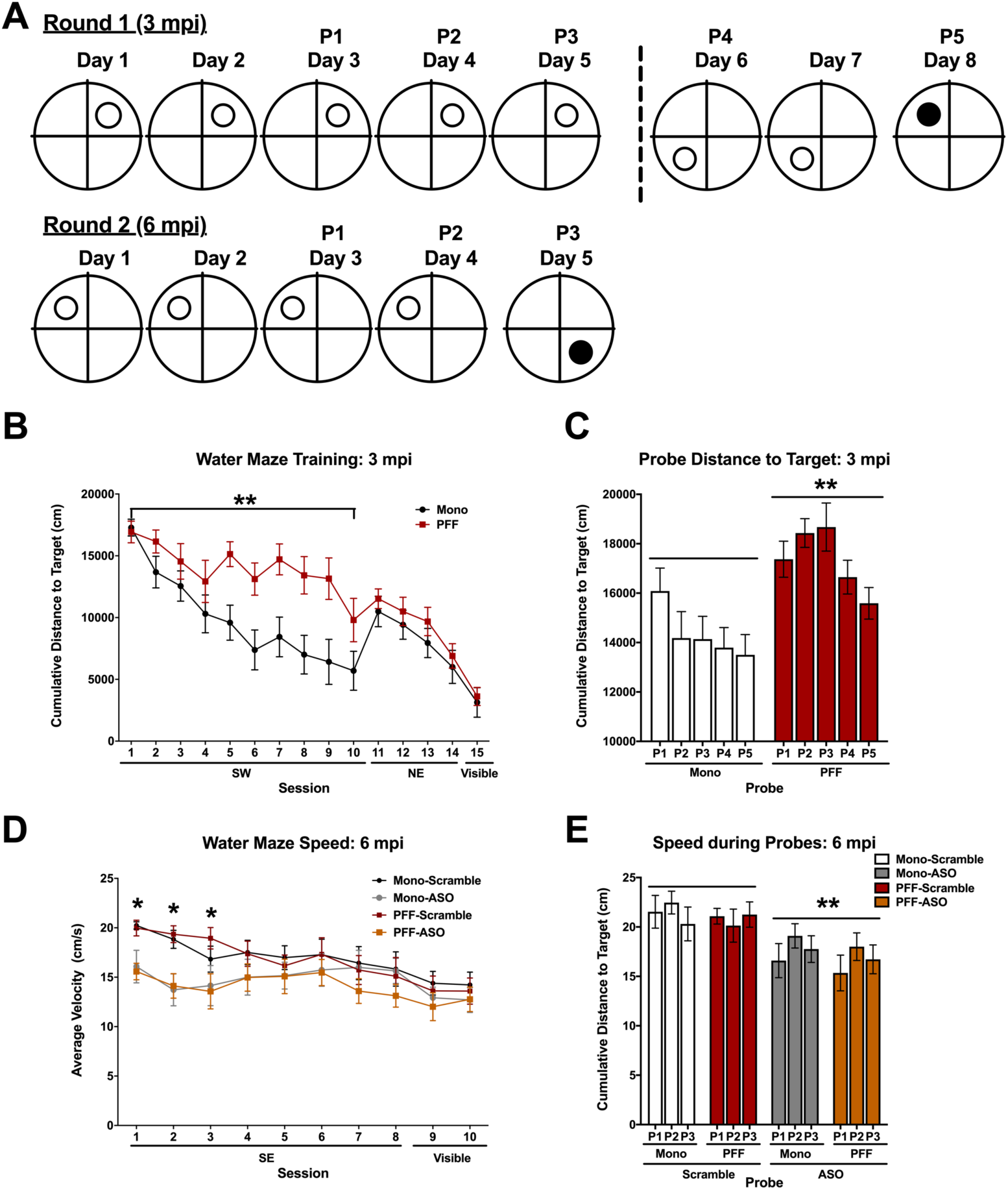
Performance in the water maze at 3 and 6 mpi. **A)** Schematic of the platform locations over the days during each round of behavioral testing. For the first round (3 mpi), the platform was located in the SW quadrant for 5 days with extra-maze cues for reference (considered “hidden” platform sessions). The platform was then moved to the NE quadrant for two days, followed by a single trial on day 8 with the extra-maze cues removed and a flag placed on the platform (considered “visible” platform sessions). Probe trials to assess memory retention were performed in the morning prior to any training sessions on days 3, 4, 5, 6, and 8. For the second round (6 pmi), the platform was located in the SE quadrant for 4 days of hidden sessions, followed by a single day of visible sessions in the NW quadrant. Probe trials were performed in the morning prior to training sessions on days 3, 4, and 5. **B)** Cumulative distance to the target platform over the course of training sessions during the first round of testing. For the first platform location, a significant difference was detected between PFF and Mono groups (main effect of PFF, *F*(1,31) = 9.373, *p* = 0.005). No difference was detected when the platform location changed, nor during the visible platform session. **C)** Cumulative distance to the target platform location (where the platform was located during the preceding hidden platform training trials) during probe trials at 3 mpi. Over all probe trials, PFF animals swam farther from the platform than Mono animals (main effect of PFF, *F*(1,31) = 11.622, *p* = 0.002). **D)** Average swim speeds during training in the second round of behavioral testing. ASO-treated animals swam slower than control-treated animals (main effect of ASO, *F*(1,29) = 5.693, *p* = 0.024). Multivariate analysis revealed significance based on ASO for the first three trials. **E)** Average swim speeds during the probe trials in the second round of behavioral testing. ASO animals swam slower than control animals (main effect of ASO, *F*(1,29) = 7.865, *p* = 0.009). Data are presented as group averages ±SEM; \**p* < 0.05, \*\**p* < 0.01.

Throughout training, we periodically assessed spatial memory retention using probe trials, during which the platform was removed. Similar to training, we saw a higher cumulative distance to the target location in PFF mice compared to Mono mice (repeated measures ANOVA: time x PFF interaction *F*(4,124) = 2.641, *p* = 0.037, sphericity assumed; main effect of PFF, *F*(1,31) = 11.622, *p* = 0.002; Fig. 2C), suggesting an impairment in spatial memory. This effect was found for all target locations. Along with that, though, PFF-injected mice swam slower during these probe trials (time x PFF interaction *F*(4,116) = 15.277, *p* = 0.036, sphericity assumed; main effect of PFF, *F*(1,29) = 6.025, *p* = 0.02; data not shown).

After ASO delivery, we performed the WM test again, with new platform locations not used in the first round of testing. When spatial learning was assessed, we discovered that mice treated with this particular ASO swam significantly slower than control animals (2-way repeated measures ANOVA: main effect of ASO, *F*(1,29) = 5.698, *p* = 0.024; Fig. 2D), but did not detect a difference in swim speeds based on PFF status. These ASO-treated mice also had slower average swim speeds during probe trials (2-way repeated measures ANOVA: time x ASO interaction, *F*(2,58) = 4.476, *p* = 0.016, sphericity assumed; main effect of ASO, *F*(1,29) = 7.865, *p =* 0.009; Fig. 2E). There were no differences in speed or distance from the target platform during training, and no difference in distance from the target during probes.

### Novel object recognition and spatial habituation are impaired in PFF mice at 6 mpi, but rescued by targeted ASO treatment

We tested animals in the open field test both pre-and post-ASO delivery to assess anxiety-like behavior as well as spatial habituation learning. At 3 mpi, there were no differences detected in total distance moved, average velocity, or time spent in the center, with both PFF-and Mono-injected animals showing typical habituation to the enclosures. At 6 mpi, though, we saw an intriguing time-by-PFF-by-ASO interactions in the total distance moved (2-way repeated measures ANOVA: time x PFF x ASO, *F*(1,29) = 5.637, *p* = 0.024, sphericity assumed; Fig. 3A) and average velocity (time x PFF x ASO, *F*(1,29) = 5.533, *p* = 0.026, sphericity assumed). All groups except for PFF-Scramble mice showed the expected decrease in overall activity and velocity, indicating spatial habituation, whereas PFF-Scramble animals showed a slight increase in both measures.

**Figure 3.**
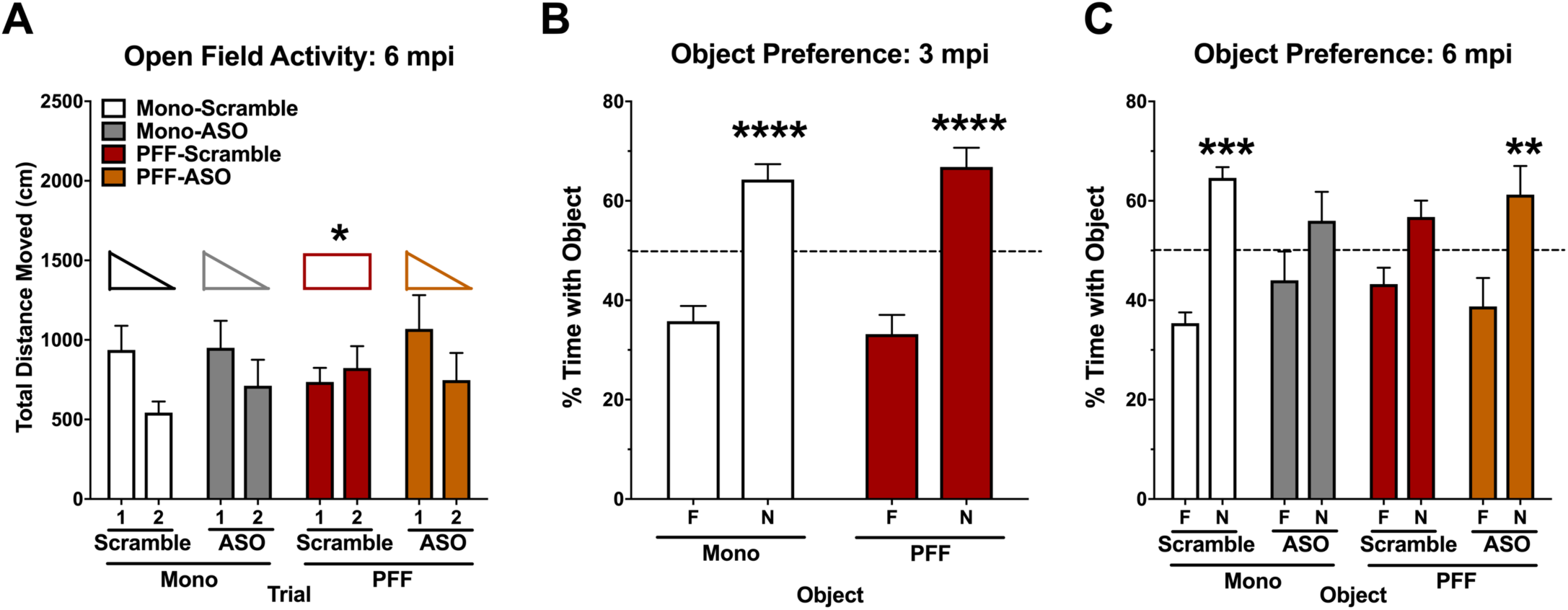
Performance in the open field and novel object recognition. **A)** Total distance moved in the open field in the second round of behavioral testing (6 mpi). PFF-Scramble animals showed an impairment in spatial habituation (time x PFF x ASO interaction, *F*(1,29) = 5.637, *p* = 0.024) whereas all other groups showed the expected decline in total distance moved on the second day. **B)** Object recognition memory on test day during the first round of behavioral testing. Both Mono-and PFF-mice showed robust preference for the novel object on test day (Sidak’s multiple comparisons, *p* < 0.0001 for both). **D)** Object recognition memory on test day during the second round of behavioral testing. Mono-Scramble and PFF-ASO animals showed a preference for the novel object *(p* = 0.0004, *p* = 0.0043, respectively), whereas Mono-ASO and PFF-Scramble animals did not *(p* = 0.2541, *p* = 0.2099, respectively). Data are presented as group averages ±SEM. Animals that explored <2s were excluded from analysis. * *p* <0.05, ** *p* <0.01, *** *p* <0.001, **** *p* <0.0001.

Following open field testing, we assessed novel object recognition in the same enclosures. Mice were first exposed to the open field containing two identical objects; 24 h later, one familiar object (F) was replaced with a distinct, novel object (N) for object recognition testing. We implemented this test at both 3 and 6 mpi. During the second round of behavior, different objects were used than during the first round. Time spent with the novel object compared to the familiar object was analyzed as a measure of memory. At 3 mpi, there were no differences seen in object memory, with both groups showing robust preference for the novel object (Sidak’s multiple comparisons; Mono familiar vs. novel, *t*(56) = 5.168, *p* < 0.0001; PFF familiar vs. novel, *t*(56) = 6.968, *p* < 0.0001; Fig. 3B). At 6 mpi, we again saw differential effects of ASO based on PFF status. Mono-Scramble and PFF-ASO animals both showed a preference to explore the novel object (Mono-Scramble familiar vs. novel, *t*(52) = 4.206, *p* = 0.0004; PFF-ASO familiar vs. novel, *t*(52) = 3.462, *p* = 0.0043; Fig. 3C). However, neither Mono-ASO nor PFF-Scramble showed a preference to explore the novel object (Mono-ASO familiar vs. novel, *t*(52) = 1.845, *p* = 0.2541; PFF-Scramble familiar vs. novel, *t*(52) = 1.945, *p* = 0.2099; Fig. 3C). These results suggest that this particular mouse ASO at this dose has a negative effect on hippocampus-dependent cognitive function, as assessed in the object preference test.

### PFF and ASO injections are associated with impairments in motor function

To assess the potential effects of PFF and ASO on motor function, we tested all animals in the rotarod and wire hang tests at two time points (3 mpi and 6 mpi; Fig. 4A & 4B). Based on the timeline of our injections, we expected to see subtle differences between PFF-and Mono-injected animals at 3 mpi and more pronounced motor impairments in PFF animals at 6 mpi.

**Figure 4.**
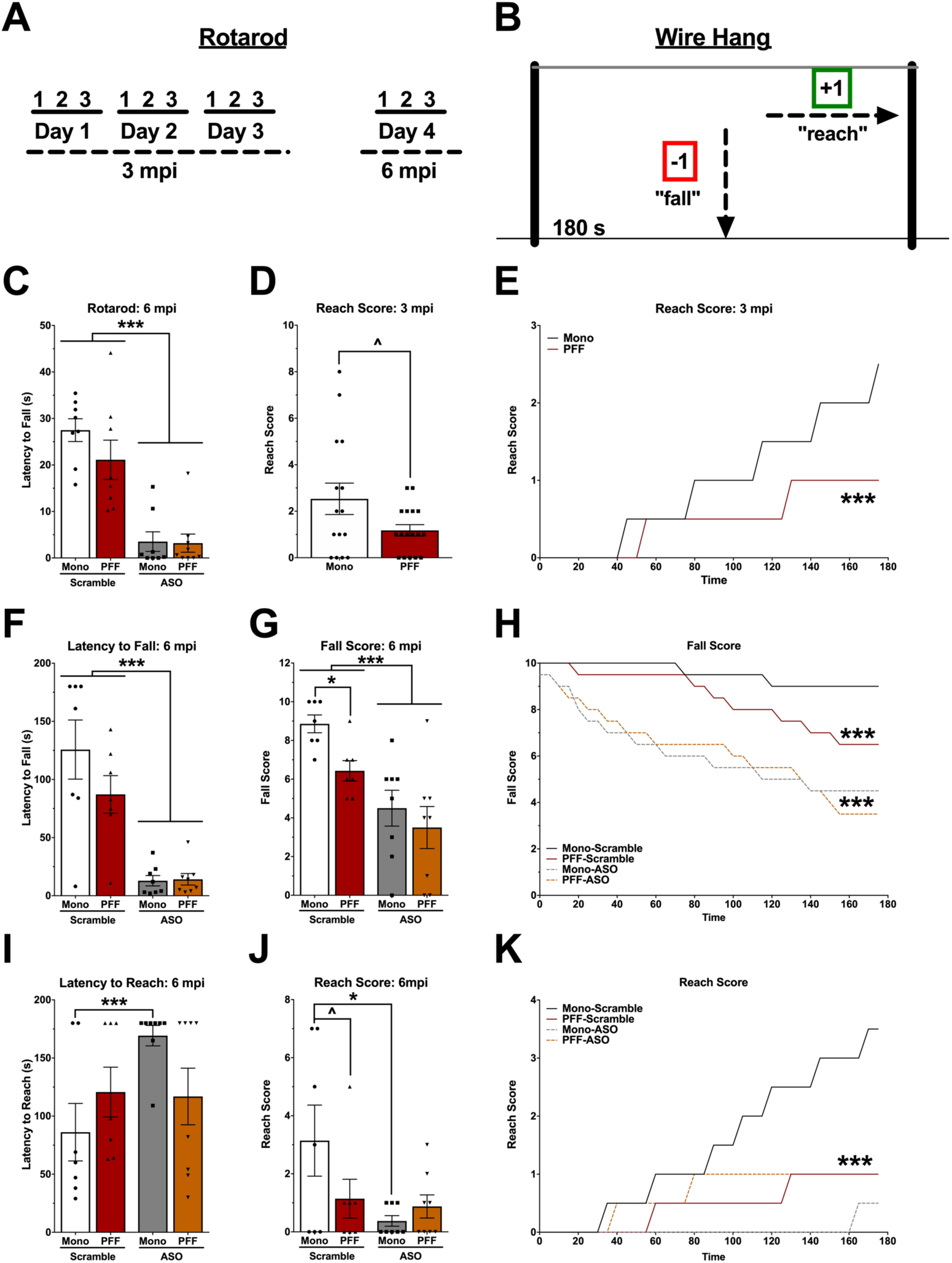
Motor Measures. **A)** Schematic of the organization of the rotarod test. At 3 mpi, animals received 3 subsequent trials each day over a 3-day period. At 6 mpi, animals received a single day of 3 subsequent trials. **B)** Schematic of the “falls and reaches” wire hang method. Method was the same for both 3 and 6 mpi. **C)** 6 mpi average latency to fall off the rotarod. A 2-way ANOVA did not show a difference based on PFF status; however, ASO animals fell significantly earlier than control animals (*F*(1,28) = 52.316, *p* < 0.001). **D)** 3 mpi reach score in the wire hang. An independent samples t-test indicated a trend towards PFF animals having a lower final score than Mono animals (*t*(30) = 1.981, *p* = 0.057). **E)** 3 mpi reach score over the 180 s wire hang trial. A repeated measures ANOVA indicated a significant PFF by time interaction (*F*(35,980) = 3.195, *p* < 0.001), where Mono mice showed a faster increase in reach score than PFF mice. **F)** 6 mpi latency to fall off the wire hang. A 2-way ANOVA showed ASO animals fell sooner than control (*F*(1,26) = 41.25, *p* < 0.0001); there was no effect of PFF. **G)** 6 mpi final fall score. ASO treated mice had a lower final fall score (*F*(1,26) = 19.14, *p* < 0.001) than controls. PFF animals also had a lower fall score than Mono animals (*F*(1,26) = 4.239, *p* < 0.05). **H)** 6 mpi fall score plotted over the 180 s. A repeated measures ANOVA revealed a time by ASO interaction (*F*(35,910) = 6.087, *p* < 0.001) and a time by PFF interaction (*F*(35,910) = 3.409, *p* < 0.001), with Mono-Scramble falling the least over time, PFF-Scramble animals falling more over time, and both ASO groups falling the most. **I)** 6 mpi latency to reach in the wire hang. A 2-way ANOVA for latency to first reach showed a PFF by ASO interaction (*F*(1,26) = 4.443, *p* < 0.05), with Mono-ASO animals performing worse than Mono-Scramble animals, but PFF animals performing the same regardless of ASO treatment. **J)** 6 mpi final reach score. Similar results were seen with final reach score, with a main effect of ASO (*F*(1,26) = 4.837, *p* < 0.05) and a trend toward a PFF by ASO interaction (*F*(1,26) = 3.218, *p* = 0.0817). Mono-ASO animals had a lower reach score than Mono-Scramble, but PFF animals did not differ in score based on ASO treatment. **K)** The average reach score plotted over 180 seconds. A repeated measures ANOVA showed a time by ASO by PFF interaction (*F*(35,910) = 4.036, *p* < 0.001), a time by ASO interaction (*F*(35,910) = 5.116, *p* < 0.001) and a PFF-by-ASO interaction (*F*(1,26) = 6.770, *p* < 0.05). Mono-Scramble animals reached more over time than PFF animals, and Mono-ASO animals almost never reached. Data are presented as group averages ±SEM; individual points are individual animals. *** *p* < 0.001, * *p* <0.05.

At 3 mpi, PFF mice did not show a decreased ability to stay on the rotarod (1-way repeated measures ANOVA; *F*(1,31) = 0.000, *p* = 0.995; Table 2) or the wire hang (*t*(32) = 0.229, *p* = 0.8205; Table 2) compared to Mono-injected mice. However, there was a trend toward a decreased final reach score in PFF compared to Mono animals (*t*(30) = 1.981, *p* = 0.057; Fig. 4D). When reach score was assessed over the 180 seconds, a significant time-by-PFF interaction was revealed (1-way repeated measures ANOVA; *F*(35,980) = 3.195, *p* < 0.001), suggesting a mild impairment in coordinated motor movement at 3 mpi (Fig. 4E).

**Table 2.** Performance in motor tasks at 3 mpi. There was a trend towards a significant effect of PFF in the wire hang reach score (p = 0.057). No other differences were detected between Mono-and PFF-injected animals. All data are presented as averages ±SEM.

| Test | Measure | Mono | PFF | Statistical Test | *p* value |
| --- | --- | --- | --- | --- | --- |
| Rotarod | Fall Latency (s) | 43.28 | 43.96 | RM-ANOVA | 0.995 |
| Wire Hang | Fall Latency (s) | 97.69 | 100.18 | t-test | 0.821 |
| Wire Hang | Fall Score | 8.13 | 8.06 | t-test | 0.498 |
| Wire Hang | Reach Latency (s) | 110.50 | 118.29 | t-test | 0.548 |

At 6 mpi, there were no differences in rotarod performance based on PFF status (2-way repeated measures ANOVA (*F*(1,28) = 1.207, *p* = 0.281; Fig. 4C). Interestingly, there was a pronounced deficit in the ability of animals treated with this particular mouse ASO to stay on the rotarod (Time x ASO interaction: *F*(2,56) = 29.496, *p* < 0.001; Main Effect of ASO: *F*(1,28) = 52.316, *p* < 0.001; Fig. 4C). In the wire hang, PFF-injected mice fell more compared to Mono controls (main effect of PFF: *F*(1,26) = 4.239, *p* < 0.05; Fig. 4G) at this time point. These ASO-treated mice also fell sooner (2-way ANOVA; main effect of ASO: *F*(1,26) = 41.25, *p* < 0.0001; Fig. 4F) and more often (main effect of ASO: *F*(1,26) = 19.14, *p* < 0.001) than their control counterparts (Fig. 4G). When fall scores were analyzed over the 180 seconds, there were significant time-by-ASO (*F*(35,910) = 6.087, *p* < 0.001) and time-by-PFF (*F*(35,910) = 3.409, *p* < 0.001) interactions, with PFF-Scramble animals performing worse than Mono-Scramble animals, and both PFF and Mono mice treated with this ASOs performing worse than both control ASO groups (Fig. 4H).

Lastly, coordinated motor movement measured on the wire hang was differentially affected, shown by a PFF-by-ASO interaction in the latency to reach the edges of the wire hang (*F*(1,26) = 4.443, *p* < 0.05; Fig. 4I). Analysis of the final reach score showed a main effect of ASO (*F*(1,26) = 4.837, *p* < 0.05) and a trend toward a PFF by ASO interaction (*F*(1,26) = 3.218, *p* = 0.0817; Fig. 4J). Mono-ASO animals had a lower reach score than Mono-Scramble, but PFF animals did not differ in score based on ASO treatment. This was also shown when analyzing the reach score over time, with a time-by-PFF-by-ASO interaction (*F*(35,910) = 4.036, *p* < 0.001), in addition to a time-by-ASO interaction (*F*(35,910) = 5.116, *p* < 0.001) and a significant PFF-by-ASO interaction (*F*(1,26) = 6.770, *p* < 0.05; Fig. 4K). Mono-Scramble animals performed the best, reaching the edges faster and more often than any other group, and Mono-ASO animals performed the worst. The PFF-Scramble and PFF-ASO mice were not different from each other in their ability to escape the wire hang.

These tests show that, in addition to the expected progression of motor impairments 6 months following PFF seeding, this particular αsyn-targeted ASO was associated with impaired motor function, which may be due to unknown off-target effects not involving ASO-mediated lowering of αsyn.

### WT and αsyn knock out mice treated with ASO show alterations in sleep patterns, feeding behavior, and weight loss, indicating possible off-target effects

Sleep disturbances are often reported in patients with LB pathology, with evidence that sleep disorders can appear nearly a decade before other symptoms emerge (Loddo et al., 2017). To determine whether there were alterations in circadian activity levels, animals were placed into home-cage monitoring devices to assess undisturbed activity over a week. Over the course of the week, mice treated with this particular ASO showed alterations in activity, specifically during the light cycle (Day x ASO interaction: *F*(1.703,34.060) = 3.550, *p* < 0.05, Huynh-Feldt corrected). While control-treated mice decreased their average daytime activity over the week, ASO-mice showed an *increase* in their average activity during the light period (Fig. 5A). This time-by-ASO interaction was not observed during nighttime activity over the week (Fig. 5B). When analyzing light and dark cycles in half hour increments, the data showed a shift in dark-to-light transitions for two of the four mornings recorded *(Tuesday,* Time x ASO interaction: *F*(18.730,65.638) = 1.719, *p* < 0.05; *Wednesday,* Time x ASO interaction: *F*(22,440) = 1.782, *p* = 0.017), with three of the four mornings trending towards a general main effect of ASO being associated with higher activity *(Wednesday,* ASO: *F*(1,20) = 3.471, *p* = 0.077; *Thursday,* ASO: *F*(1,20) = 3.328, *p* =0.083; *Friday,* ASO: *F*(1,20) = 2.996, *p* = 0.099). ASO mice appeared to have a delayed sleep onset, as their activity extended past lights-on longer than that of control-treated mice. A similar shift in sleep/wake transition was not observed at the light-to-dark transition for any of the nights recorded.

**Figure 5.**
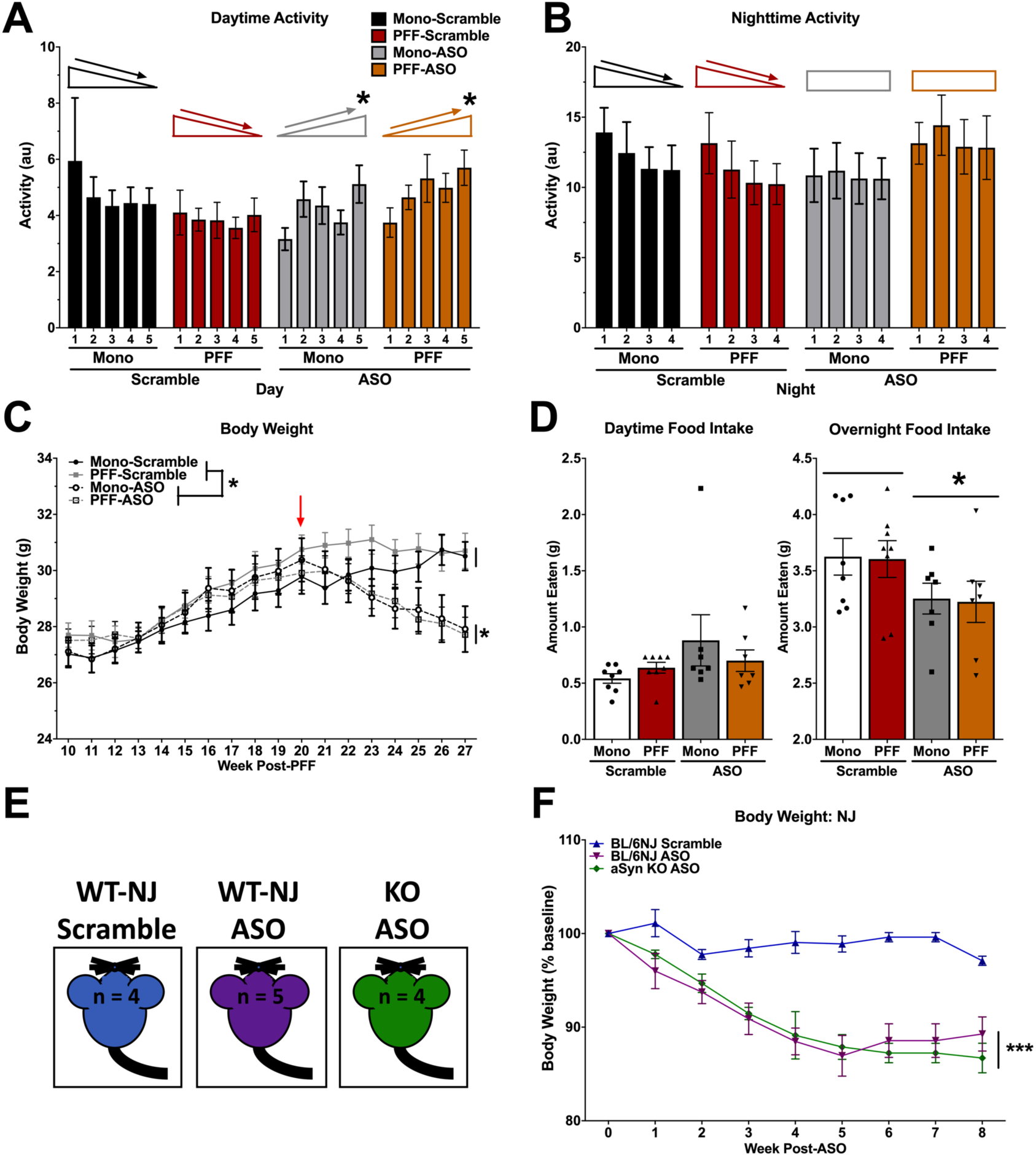
Health-related data. **A)** Average daytime activity over the course of 5 days. A repeated measures ANOVA for daytime activity indicated an interaction of day and ASO (*F*(1.703,34.060) = 3.550, *p* < 0.05), where control animals decreased their daytime activity over the week, but ASO animals increased. **B)** Average nighttime activity over the course of 6 nights. A repeated measures ANOVA for nighttime activity suggested a similar pattern. Control animals showed a significant decrease in activity over nights (Effect of night, *F*(1.848,18.477) = 7.529, *p* = 0.005), whereas ASO animals did not (no effect of night, *F*(3,30) = 0.57, *p* = 0.639). **C)** Body weights were recorded weekly beginning at the first round of behavioral testing. Prior to ASO delivery, a repeated-measures ANOVA showed that all animals increased in body weight over time (*F*(3.229,96.862) = 122.880, *p* < 0.001) with no group differences in weight. Following ASO delivery, ASO animals showed a steady decline in body weight over time (Time x ASO interaction: *F*(3.613,104.777) = 16.093, *p* < 0.001), and an overall reduced body weight compared to control animals (Main Effect of ASO: *F*(1,29) = 7.952, *p* < 0.001). **D)** Food intake was recorded over a 5-day period. Food was weighed in the morning (∼8:00 AM) and the evening (∼17:00 PM) to approximate nighttime and daytime food intake. A 2-way ANOVA revealed no group differences in daytime food intake, but did show a main effect of ASO for nighttime food intake (*F*(1,26) = 5.297, *p* < 0.05). **E)** Schematic of the control experiment to assess specificity of the ASO. C57Bl/6NJ (WT-NJ) and αsyn knock out mice (KO) were injected with the ASO construct, and compared against WT-NJ mice injected with control. **F)** A repeated measures ANOVA indicated a significant time by ASO interaction (*F*(16,88) = 6.079, *p* = 0.0001) and main effect of ASO (*F*(2,11) = 19.03, *p* = 0.0003), suggesting off-target effects. Data presented as group averages ±SEM; individual points are individual animals. * *p* <0.05, *** *p* <0.001.

Body weights were recorded once a week over the duration of the experiment. Prior to ASO-injections, there were no differences observed in body weight (2-way repeated measures ANOVA, *F*(1,30) = 1.203, *p* = 0.281), with all groups showing an expected increase over time (Time: *F*(3.229,96.862) = 122.880, *p* < 0.001). However, following ICV delivery of ASO or control, ASO-treated mice showed a steady decline in body weight (Time x ASO interaction: *F*(3.613,104.777) = 16.093, *p* < 0.001, Huynh-Feldt corrected; Main Effect of ASO: *F*(1,29) = 7.952, *p* < 0.001) until the end of the experiment (Fig. 5C). There were no differences based on PFF status (p = 0.544), nor an interaction of PFF and ASO (p = 0.517).

As we observed body weight changes following injections of ASO, we next measured food intake over a period of 5 days. A 2-way ANOVA of average food intake revealed that ASO-injected animals ate significantly less between 17:00 and 8:00 (“overnight”) than control-injected animals (*F*(1,26) = 5.297, *p* < 0.05), when mice typically eat most of their food. The average food intake from 8:00 to 17:00 (“daytime”) was not different between the groups (Fig. 5D).

To test whether this ASO could have off-target effects, we injected it into αsyn-KO mice and C57Bl/6NJ (WT-NJ) matched controls (Fig. 5E). Following ICV delivery, both the αsyn-KO mice and WT-NJ mice that received ASO progressively lost body weight compared to the control-injected mice (Time-by-ASO interaction: *F*(16,88) = 6.079, *p* = 0.0001; Main Effect of ASO: *F*(2,11) = 19.03, *p* = 0.0003; Fig. 5F). This suggests that there are αsyn-independent off-target effects with this particular ASO.

### ASO reduces alpha-and phosphorylated-synuclein protein levels and LB-like pathology, which correlates with cognitive performance

To determine the extent to which ASO treatment reduced αsyn and phosphorylated αsyn (psyn) protein levels, we used Western blot analysis of hippocampal tissues. Neural tissue was collected after the second round of behavioral testing, 6 months post-PFF or-Mono injections and 7 weeks after ICV delivery of targeted ASO or scrambled, control ASO (Fig. 1). Both αsyn and psyn forms of the protein were decreased by greater than 50% in hippocampal tissue by the ASO treatment. A two-way ANOVA for αsyn integrated density normalized to total protein revealed a main effect of ASO (*F*(1,29) = 74.33, *p* < 0.0001; Fig. 6A). Sidak’s post hoc comparison showed ASO treatment differences within Mono and PFF groups (Mono: *p* < 0.0001; PFF: *p* < 0.0001). Similarly, there was a main effect of ASO for psyn integrated density (*F*(1,29) = 35.88, *p* < 0.0001; Fig. 6A), with Sidak’s post hoc revealing treatment differences (Mono: *p* < 0.01; PFF: *p* < 0.001). There were no effects of PFF on αsyn or psyn protein levels, nor any PFF*ASO interactions, suggesting the PFF treatment did not have an effect on overall protein levels. Representative blots are presented in Fig. 6B.

**Figure 6.**
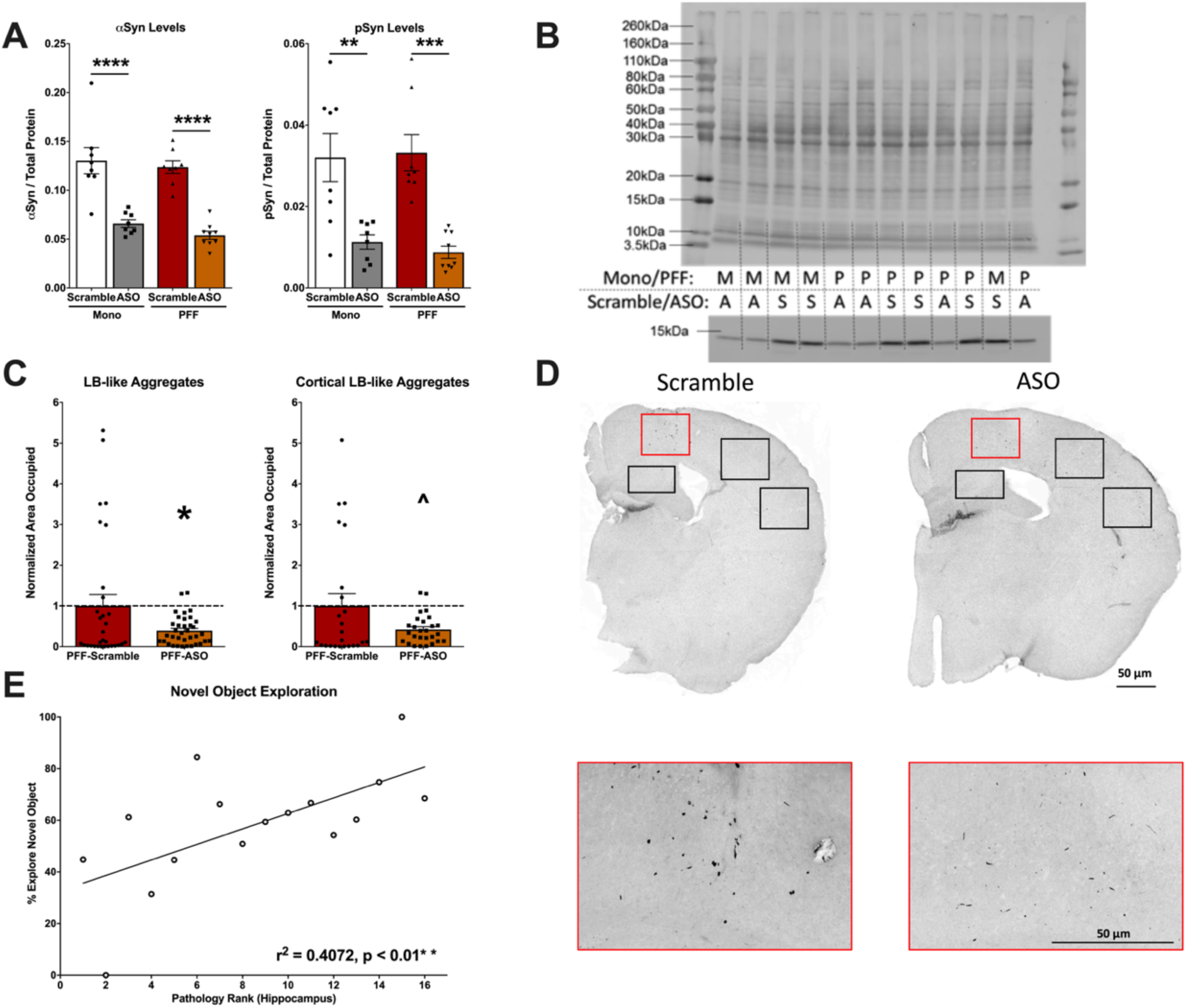
Analysis of α-synuclein (αsyn) and phosphorylated synuclein (psyn) protein levels in the hippocampus and LB-like pathology. **A)** *Left:* Quantification of αsyn, normalized to total protein. A two-way ANOVA indicated a main effect of ASO treatment (*F*(1,29) = 74.33, *p* < 0.0001) and Sidak’s post-hoc tests indicated αsyn protein was decreased in Mono-ASO mice compared to Mono-Scramble (p < 0.0001) and PFF-ASO compared to PFF-Scramble (p < 0.0001). *Right:* Quantification of psyn, normalized to total protein levels. A two-way ANOVA indicated a main effect of ASO treatment (*F*(1,29) = 35.88, *p* < 0.0001) and Tukey’s post-hoc tests indicated psyn protein was decreased in Mono-ASO mice compared to Mono-Scramble (p < 0.01) and PFF-ASO compared to PFF-Scramble (p < 0.001). Data presented as group averages ±SEM; individual points are individual animals. **B)** Representative blot of total protein stain (top) and αsyn band at 14.4 kDa (bottom). Lanes are indicated with “M” for Mono, “P” for PFF, “A” for ASO, and “S” for scramble control. **C)** Normalized burden load of pSyn signal in PFF-injected animals. Percent area covered by pSyn immunoreactvity was analyzed in the motor cortex, medial somatosensory cortex, lateral somatosensory cortex, and hippocampus. For each region, pSyn immunoreactivity was normalized to the average of the PFF-Scramble group. A composite load was created to assess overall pathology burden, and an independent-samples t-test showed an overall decrease in pathology load in the PFF-ASO group compared to the PFF-Scramble group *(left*; *t*(65) = 2.280, *p* = 0.0259). When only cortical pathology load was assessed, an independent-samples t-test showed a trend towards a decrease in pathology load in the ASO group *(right*; *t*(49) = 1.975, *p* = 0.0540). **D)** Representative sections from PFF-Scramble and PFF-ASO groups. Boxes indicate the outline of the regions for percent-area occupied analysis. The red box indicates the motor cortex, which is enlarged in below. **E)** Correlations of percent area occupied in the hippocampus to time spent with the novel object. Mice were ranked 1-17, with 1 having the most pSyn immunoreactivity in the hippocampus to 17 having the least. Percent time with the novel object was compared with a linear regression, and revealed a significant positive correlation, where animals with the most pathology spent the least amount of time with the novel object (r^2^ = 0.4072, *p* < 0.01). When mice were ranked within each group and compared with a linear regression, a significant positive correlation was shown in the PFF-Scramble group (r^2^ = 0.5746, *p* < 0.05), and a trend in the PFF-ASO group (r^2^ = 0.3894, *p* = 0.0725). * *p* <0.05, ** *p* < 0.01, *** *p* < 0.001.

Next, we assessed if ASO reduced the LB-like pathology load in the PFF animals using immunohistochemistry to examine psyn-positive aggregates. We analyzed the percent area occupied by LB-like pathology in four different brain regions (motor cortex, medial somatosensory cortex, lateral somatosensory cortex, and hippocampus) to create a composite pathology burden measure. Using this composite score, we detected a significant reduction in LB-like aggregate load in ASO-treated mice compared to control (*t*(65) = 2.280, *p* = 0.0259; Fig. 6C, left). When only cortical regions were analyzed, we observed a trend toward a significant difference (*t*(49) = 1.975, *p* = 0.054; Fig. 6C, right). However, when we analyzed each individual region separately, we did not observe significant differences between PFF animals treated with ASO or control (Table 3; t-tests; motor cortex, *p* = 0.2817; medial somatosensory cortex, *p* = 0.2701; lateral somatosensory cortex, *p* = 0.323; hippocampus, *p* = 0.3039). Representative images of ASO-and control-treated PFF hemispheres are shown in Fig. 6D.

**Table 3.**
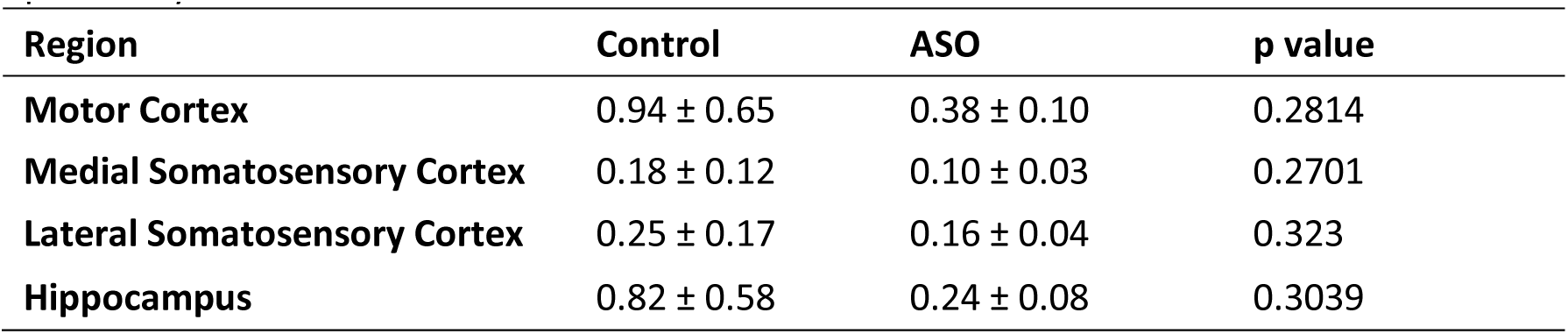
pSyn immunoreactivity in each brain region. Data are the average percent area occupied ±SEM. *p* values were calculated using independent samples t-tests, and did not reveal any significant differences when each region was analyzed independently.

We next assessed whether the pathology burden was related to the behavioral and cognitive phenotypes. Thus, we ran correlational analyses to determine if amount of pathology corresponded to our observed behavioral alterations. A linear regression of pathology load in the motor cortex did not reveal any significant correlations with any of the motor behavioral measures. Interestingly, a linear regression of pathology in the hippocampus significantly inversely correlated with percent time exploring the novel object in the second round of NOR testing *(r^2^* = 0.4072, *p* < 0.01). Mice with more LB-like pathology explored the novel object less (Fig. 6E). When split into groups based on ASO-or control-injection, linear regressions of each showed that this correlation held up in PFF-Scramble animals *(r^2^* = 0.5746, *p* < 0.05), but was attenuated by ASO injection *(r*^2^ = 0.3894, *p* = 0.0725).

## Discussion

We set out to assess the progressive cognitive and behavioral effects of PFF injections in the motor cortex of male WT, as well as explore if reducing αsyn expression throughout the CNS with a targeted ASO could ameliorate cognitive, behavioral, and pathological effects of PFF injections. The PFF model of synucleinopathies has become more common and is gaining in popularity since it was first described (Luk, Kehm, Carroll, et al., 2012; Volpicelli-Daley et al., 2011). Our lab has previously described the pathological progression of motor cortex injections using in-vivo multiphoton imaging (Osterberg et al., 2015) but no one has characterized the phenotypic progression following motor cortex injections. The results of the current study provide supporting evidence that there is a progressive decline in behavioral and cognitive performance following PFF injections, as PFF-injected mice showed mild motor and cognitive impairments at 3 months post-injection (mpi) that became more pronounced at 6 mpi. Progression of motor impairment was particularly pronounced, as PFF-injected mice performed similarly to Mono-injected controls in the wire hang at 3 mpi, but fell sooner and more often and were unable to “escape” at similar rates at 6 mpi. Clinically, the motor deficits observed in PD are likely reflective of a combination of problems, including bradykinesia and rigidity-which result in the difficulty to execute motor tasks, especially repetitive tasks-as escaping from the wire hang requires fine, repetitive motor control of all four paws (Berardelli, Rothwell, Thompson, & Hallett, 2001; Giovannoni, van Schalkwyk, Fritz, & Lees, 1999; Jankovic, 2008; van Putten, 2016). These results are similar to results from other groups that have assessed motor performance following striatal injections: original characterization of striatal injections indicated that wire hang deficits began at 3 mpi, while rotarod deficits did not appear until 6 mpi and locomotion in the open field remained unaffected up to 6 mpi (Luk, Kehm, Carroll, et al., 2012). Minor deficits in the balance beam have been reported 2 months post-striatal injection without differences in rotarod or open field locomotion, as well (Karampetsou et al., 2017).

Additionally, we observed hippocampus-dependent cognitive impairment at both 3 mpi and 6 mpi, suggesting spread of pathology from the motor cortex to the hippocampus. The hippocampus has only recently become a region of interest in PD, with evidence showing that disrupted dopaminergic signaling impairs hippocampal plasticity (Calabresi, Castrioto, Di Filippo, & Picconi, 2013). Dopamine signaling from the midbrain to the hippocampus has been shown in both rodents and humans to modulate long-term learning. Healthy adults who were placed in an MRI during an image-recognition task showed increased BOLD signal in the midbrain and hippocampus in response to previously-seen, reward-associated images (Wittmann et al., 2005). In rodents, both a toxin-induced model and a transgenic model of PD showed impaired long-term potentiation in the hippocampus that was associated with decreased dopamine transmission and impaired hippocampus-dependent spatial memory (Costa et al., 2012). Here, we saw hippocampus-dependent novel object recognition in PFF animals at 3 mpi, but not at 6 mpi; Mono-injected mice showed object preference at both time points. Post-mortem analysis of LB-like pathology confirmed that psyn-positive inclusions had spread into the hippocampus, and amount of pathology inversely correlated with novel object performance. At 3 mpi, PFF mice showed learning and memory impairments in the water maze, measured by the higher cumulative distance to the target platform both during hidden platform training and probe trails. No impairment was seen when the platform location was switched at 3 mpi, nor at 6 mpi, which could be due to an over-training effect eliminating the ability to detect differences. An over-training effect can eliminate differences even in rats that have had hippocampal and/or subiculum lesions (Morris, Schenk, Tweedie, & Jarrard, 1990). Others have demonstrated how over training can hide learning and memory differences in an age-dependent manner in an Alzheimer’s mouse model, also (Weitzner, Engler-Chiurazzi, Kotilinek, Ashe, & Reed, 2015).

Assessment of the αsyn-targeted ASO indicated rescue of some behavioral and cognitive impairments, and decreased LB-like pathology, but also revealed possible off-target effects. The ASO construct we used reduced hippocampal αsyn and psyn by >50%, and successfully ameliorated the amount of psyn-positive inclusions. Presumably this reduction in LB-like pathology occurred after LB-like inclusions had begun forming, supporting this approach’s potential in the clinic, where patients begin treatment well after LBs have formed and spread to many brain regions (Braak et al., 2003; Del Rey et al., 2018). Our timing for ASO administration was aimed at reflecting the timing that is typical of diagnosis of synucleinopathies in the clinic, with a goal to guarantee robust αsyn reduction. Here, ASO treatment was associated with improved performance in the novel object recognition test: while PFF-scramble animals did not show a preference, PFF-ASO animals demonstrated intact hippocampus-dependent object recognition. Additionally, PFF-ASO mice displayed typical spatial habituation in the open field, which was absent in PFF-Scramble animals. Others have reported evidence that ASOs could rescue disrupted dopamine signaling: administration of a distinct αsyn-specific ASO to WT mice, which resulted in 20-40% decrease of αsyn, increased dopamine and serotonin release in the forebrain, whereas αsyn over-expression decreased signaling (Alarcon-Aris et al., 2018). While we did not assess dopaminergic signaling, it is possible that the ASO construct similarly maintained dopaminergic signaling to the hippocampus, resulting in rescued hippocampus-dependent memory. These results suggest that the ASO had beneficial effects in the context of disease and could be a potential treatment to reduce pathological spread.

While these results indicated that reducing αsyn could be beneficial in a diseased state, we also observed some detrimental side effects on behavioral measures using this particular mouse ASO, including disruptions in light-dark cycles, body weight, food intake, and motor abilities. Noticeably, the effects on general health measures were significant, with ASO-treated animals showing declining body weight, lower food intake, and delayed onset of sleep regardless of PFF status. Another assessment of an αsyn-specific ASO did not report changes in general health, but a lower dose (300μg) was administered in this case (Alarcon-Aris et al., 2018). Our ASO dose (700 μg) caused a more substantial reduction (>50%) than this previous study (20-40%), which could account for the differential results. Importantly, though, similar weight loss was also seen in αsyn knock out animals, indicating that these results are most likely due to off-target effects of our particular ASO construct, and not necessarily attributed to ASO-mediated lowering of αsyn.

The most profound motor deficits we observed were in the wire hang and rotarod tests, with ASO-treated mice falling significantly more in both tasks and showing impaired motor coordination as measured by their ability to escape the wire hang. PFF-injected mice also showed a decreased number of escapes from the wire hang and a higher latency to the first escape: in this circumstance, our ASO delivery did not further worsen the performance of the PFF mice, but did significantly impair the Mono mice, suggesting that the unknown off-target effects had similar negative consequences.

While our results suggest that the particular ASO we used had off-target effects, discussions about the normal role of soluble αsyn are important to consider as well. There are few tolerability concerns reported for constitutive genetic *SNCA* deletion in *SNCA* knockout mice (Abeliovich et al., 2000; Cabin et al., 2002; Chandra et al., 2004; Goldberg and Lansbury, 2000; Greten-Harrison et al., 2010). The effects of reducing αsyn in adult animals has produced somewhat conflicting results. RNAi-mediated αsyn silencing in the substantia nigra of rats led to degeneration of dopamine cells and neuroinflammation (Benskey et al., 2018; Gorbatyuk et al., 2010; Khodr et al., 2011). However, other groups have shown that knockdown of αsyn in the substantia nigra of adult rats using an RNAi was long-lasting and safe, with significant reduction of αsyn and no neurodegeneration observed 12 months later (Zharikov et al., 2019). In addition, multiple other groups using siRNAs, shRNAs, or ASOs against *SCNA* in mice report no cell death *in vivo* (Cole et al., 2019; Alarcon-Aris et al., 2018; Uehara et al., 2019; Zharikov et al., 2015). Two αsyn antibodies, PRX002 and BIIB054, completed first-in-human trials in which no serious adverse events were found with αsyn lowering (Schenk et al., 2017, Brys et al., 2019). It will be important in future studies to better define the physiological and functional importance of soluble αsyn.

In summary, our data suggest that αsyn reduction has beneficial consequences, and can successfully reduce LB-like pathology and rescue behavioral and cognitive deterioration. Our results support a promising future for the use of ASOs in the clinic. Ongoing clinical trials and future efforts with more advanced target-specific ASOs are warranted for determining the extent to which targeted ASO treatment can slow or halt disease progression.

## Acknowledgements

The authors would like to thank those who contributed by assisting with data acquisition and analysis (Eileen Ruth Torres, Joanne S. Lee, Crystal Chaw, Stefanie Kaech Petrie, Esha M. Patel), teaching techniques (Sydney Elise Dent, Allison J. Schaser, Teresa Stackhouse, Valerie Osterberg), and supplying reagents (Hien Tran Zhao, Kelvin Luk).

## References

Alarcon-Aris, D., Recasens, A., Galofre, M., Carballo-Carbajal, I., Zacchi, N., Ruiz-Bronchal, E., … Bortolozzi, A. (2018). Selective alpha-Synuclein Knockdown in Monoamine Neurons by Intranasal Oligonucleotide Delivery: Potential Therapy for Parkinson’s Disease. Mol Ther, 26(2), 550–567. doi:10.1016/j.ymthe.2017.11.015

Ansari, K. A., & Johnson, A. (1975). Olfactory function in patients with Parkinson’s disease. J Chronic Dis, 28(9), 493–497. doi:10.1016/0021-9681(75)90058-2

Appel-Cresswell, S., Vilarino-Guell, C., Encarnacion, M., Sherman, H., Yu, I., Shah, B., … Farrer, M. J. (2013). Alpha-synuclein p.H50Q, a novel pathogenic mutation for Parkinson’s disease. Mov Disord, 28(6), 811–813. doi:10.1002/mds.25421

Benskey, M. J., Sellnow, R. C., Sandoval, I. M., Sortwell, C. E., Lipton, J. W., & Manfredsson, F. P. (2018). Silencing Alpha Synuclein in Mature Nigral Neurons Results in Rapid Neuroinflammation and Subsequent Toxicity. Front Mol Neurosci, 11, 36. doi:10.3389/fnmol.2018.00036

Berardelli, A., Rothwell, J. C., Thompson, P. D., & Hallett, M. (2001). Pathophysiology of bradykinesia in Parkinson’s disease. Brain, 124*(Pt* 11), 2131–2146. doi:10.1093/brain/124.11.2131

Bocchio, M., Nabavi, S., & Capogna, M. (2017). Synaptic Plasticity, Engrams, and Network Oscillations in Amygdala Circuits for Storage and Retrieval of Emotional Memories. Neuron, 94(4), 731–743. doi:10.1016/j.neuron.2017.03.022

Braak, H., Del Tredici, K., Rub, U., de Vos, R. A., Jansen Steur, E. N., & Braak, E. (2003). Staging of brain pathology related to sporadic Parkinson’s disease. Neurobiol Aging, 24(2), 197–211. doi:10.1016/s0197-4580(02)00065-9

Brettschneider, J., Del Tredici, K., Lee, V. M., & Trojanowski, J. Q. (2015). Spreading of pathology in neurodegenerative diseases: a focus on human studies. Nat Rev Neurosci, 16(2), 109–120. doi:10.1038/nrn3887

Brundin, P., Ma, J., & Kordower, J. H. (2016). How strong is the evidence that Parkinson’s disease is a prion disorder? Curr Opin Neurol, 29(4), 459–466. doi:10.1097/WCO.0000000000000349

Brundin, P., & Melki, R. (2017). Prying into the Prion Hypothesis for Parkinson’s Disease. J Neurosci, 37(41), 9808–9818. doi:10.1523/JNEUROSCI.1788-16.2017

Burre, J., Sharma, M., & Sudhof, T. C. (2014). alpha-Synuclein assembles into higher-order multimers upon membrane binding to promote SNARE complex formation. Proc Natl Acad Sci USA, 111(40), E4274–4283. doi:10.1073/pnas.1416598111

Burre, J., Sharma, M., Tsetsenis, T., Buchman, V., Etherton, M. R., & Sudhof, T. C. (2010). Alpha-synuclein promotes SNARE-complex assembly in vivo and in vitro. Science, 329(5999), 1663–1667. doi:10.1126/science.1195227

Calabresi, P., Castrioto, A., Di Filippo, M., & Picconi, B. (2013). New experimental and clinical links between the hippocampus and the dopaminergic system in Parkinson’s disease. Lancet Neurol, 12(8), 811–821. doi:10.1016/S1474-4422(13)70118-2

Chaudhuri, K. R., Healy, D. G., Schapira, A. H., & National Institute for Clinical, E. (2006). Non-motor symptoms of Parkinson’s disease: diagnosis and management. Lancet Neurol, 5(3), 235–245. doi:10.1016/S1474-4422(06)70373-8

Costa, C., Sgobio, C., Siliquini, S., Tozzi, A., Tantucci, M., Ghiglieri, V., … Calabresi, P. (2012). Mechanisms underlying the impairment of hippocampal long-term potentiation and memory in experimental Parkinson’s disease. Brain, 135(Pt 6), 1884–1899. doi:10.1093/brain/aws101

Curzon, P., Rustay, N. R., & Browman, K. E. (2009). Cued and Contextual Fear Conditioning for Rodents. In nd & J. J. Buccafusco (Eds.), Methods of Behavior Analysis in Neuroscience. Boca Raton (FL).

Del Rey, N. L., Quiroga-Varela, A., Garbayo, E., Carballo-Carbajal, I., Fernandez-Santiago, R., Monje, M. H. G., … Blesa, J. (2018). Advances in Parkinson’s Disease: 200 Years Later. Front Neuroanat, 12, 113. doi:10.3389/fnana.2018.00113

Evers, M. M., Toonen, L. J., & van Roon-Mom, W. M. (2015). Antisense oligonucleotides in therapy for neurodegenerative disorders. Adv Drug Deliv Rev, 87, 90–103. doi:10.1016/j.addr.2015.03.008

Fujioka, S., Ogaki, K., Tacik, P. M., Uitti, R. J., Ross, O. A., & Wszolek, Z. K. (2014). Update on novel familial forms of Parkinson’s disease and multiple system atrophy. Parkinsonism Relat Disord*, 20 Suppl 1,* S29-34. doi:10.1016/S1353-8020(13)70010-5

Geng, X., Lou, H., Wang, J., Li, L., Swanson, A. L., Sun, M., … Drain, P. (2011). alpha-Synuclein binds the K(ATP) channel at insulin-secretory granules and inhibits insulin secretion. Am J Physiol Endocrinol Metab, 300(2), E276–286. doi:10.1152/ajpendo.00262.2010

Giovannoni, G., van Schalkwyk, J., Fritz, V. U., & Lees, A. J. (1999). Bradykinesia akinesia inco-ordination test (BRAIN TEST): an objective computerised assessment of upper limb motor function. J Neurol Neurosurg Psychiatry, 67(5), 624–629. doi:10.1136/jnnp.67.5.624

Gorbatyuk, O. S., Li, S., Nash, K., Gorbatyuk, M., Lewin, A. S., Sullivan, L. F., … Muzyczka, N. (2010). In vivo RNAi-mediated alpha-synuclein silencing induces nigrostriatal degeneration. Mol Ther, 18(8), 1450–1457. doi:10.1038/mt.2010.115

Huang, M., Wang, B., Li, X., Fu, C., Wang, C., & Kang, X. (2019). alpha-Synuclein: A Multifunctional Player in Exocytosis, Endocytosis, and Vesicle Recycling. Front Neurosci, 13, 28. doi:10.3389/fnins.2019.00028

Jankovic, J. (2008). Parkinson’s disease: clinical features and diagnosis. J Neurol Neurosurg Psychiatry, 79(4), 368–376. doi:10.1136/jnnp.2007.131045

Jensen, P. H., Nielsen, M. S., Jakes, R., Dotti, C. G., & Goedert, M. (1998). Binding of alpha-synuclein to brain vesicles is abolished by familial Parkinson’s disease mutation. J Biol Chem, 273(41), 26292–26294. doi:10.1074/jbc.273.41.26292

Jin, H., Kanthasamy, A., Ghosh, A., Yang, Y., Anantharam, V., & Kanthasamy, A. G. (2011). alpha-Synuclein negatively regulates protein kinase Cdelta expression to suppress apoptosis in dopaminergic neurons by reducing p300 histone acetyltransferase activity. J Neurosci, 31(6), 2035–2051. doi:10.1523/JNEUROSCI.5634-10.2011

Johnson, L. A., Olsen, R. H., Merkens, L. S., DeBarber, A., Steiner, R. D., Sullivan, P. M., … Raber, J. (2014). Apolipoprotein E-low density lipoprotein receptor interaction affects spatial memory retention and brain ApoE levels in an isoform-dependent manner. Neurobiol Dis, 64, 150–162. doi:10.1016/j.nbd.2013.12.016

Kam, T. I., Mao, X., Park, H., Chou, S. C., Karuppagounder, S. S., Umanah, G. E., … Dawson, V. L. (2018). Poly(ADP-ribose) drives pathologic alpha-synuclein neurodegeneration in Parkinson’s disease. Science, 362(6414). doi:10.1126/science.aat8407

Karampetsou, M., Ardah, M. T., Semitekolou, M., Polissidis, A., Samiotaki, M., Kalomoiri, M., … Vekrellis, K. (2017). Phosphorylated exogenous alpha-synuclein fibrils exacerbate pathology and induce neuronal dysfunction in mice. Sci Rep, 7(1), 16533. doi:10.1038/s41598-017-15813-8

Khodr, C. E., Sapru, M. K., Pedapati, J., Han, Y., West, N. C., Kells, A. P., … Bohn, M. C. (2011). An alpha-synuclein AAV gene silencing vector ameliorates a behavioral deficit in a rat model of Parkinson’s disease, but displays toxicity in dopamine neurons. Brain Res, 1395, 94–107. doi:10.1016/j.brainres.2011.04.036

Kontopoulos, E., Parvin, J. D., & Feany, M. B. (2006). Alpha-synuclein acts in the nucleus to inhibit histone acetylation and promote neurotoxicity. Hum Mol Genet, 15(20), 3012–3023. doi:10.1093/hmg/ddl243

Kruger, R., Kuhn, W., Muller, T., Woitalla, D., Graeber, M., Kosel, S., … Riess, O. (1998). Ala30Pro mutation in the gene encoding alpha-synuclein in Parkinson’s disease. Nat Genet, 18(2), 106–108. doi:10.1038/ng0298-106

Lee, B. R., & Kamitani, T. (2011). Improved immunodetection of endogenous alpha-synuclein. PLoS One, 6(8), e23939. doi:10.1371/journal.pone.0023939

Lemkau, L. R., Comellas, G., Kloepper, K. D., Woods, W. S., George, J. M., & Rienstra, C. M. (2012). Mutant protein A30P alpha-synuclein adopts wild-type fibril structure, despite slower fibrillation kinetics. J Biol Chem, 287(14), 11526-11532. doi:10.1074/jbc.M111.306902

Lesage, S., Anheim, M., Letournel, F., Bousset, L., Honore, A., Rozas, N., … French Parkinson’s Disease Genetics Study, G. (2013). G51D alpha-synuclein mutation causes a novel parkinsonian-pyramidal syndrome. Ann Neurol, 73(4), 459–471. doi:10.1002/ana.23894

Liu, X., Lee, Y. J., Liou, L. C., Ren, Q., Zhang, Z., Wang, S., & Witt, S. N. (2011). Alpha-synuclein functions in the nucleus to protect against hydroxyurea-induced replication stress in yeast. Hum Mol Genet, 20(17), 3401–3414. doi:10.1093/hmg/ddr246

Loddo, G., Calandra-Buonaura, G., Sambati, L., Giannini, G., Cecere, A., Cortelli, P., & Provini, F. (2017). The Treatment of Sleep Disorders in Parkinson’s Disease: From Research to Clinical Practice. Front Neurol, 8, 42. doi:10.3389/fneur.2017.00042

Luk, K. C., Kehm, V., Carroll, J., Zhang, B., O’Brien, P., Trojanowski, J. Q., & Lee, V. M. (2012a). Pathological alpha-synuclein transmission initiates Parkinson-like neurodegeneration in nontransgenic mice. Science, 338(6109), 949–953. doi:10.1126/science.1227157

Luk, K. C., Kehm, V. M., Zhang, B., O’Brien, P., Trojanowski, J. Q., & Lee, V. M. (2012b). Intracerebral inoculation of pathological alpha-synuclein initiates a rapidly progressive neurodegenerative alpha-synucleinopathy in mice. J Exp Med, 209(5), 975–986. doi:10.1084/jem.20112457

Maroteaux, L., Campanelli, J. T., & Scheller, R. H. (1988). Synuclein: a neuron-specific protein localized to the nucleus and presynaptic nerve terminal. J Neurosci, 8(8), 2804–2815. Retrieved from https://www.ncbi.nlm.nih.gov/pubmed/3411354

McGinnis, G. J., Friedman, D., Young, K. H., Torres, E. R., Thomas, C. R., Jr., Gough, M. J., & Raber, J. (2017). Neuroinflammatory and cognitive consequences of combined radiation and immunotherapy in a novel preclinical model. Oncotarget, 8(6), 9155–9173. doi:10.18632/oncotarget.13551

Miller, I. N., & Cronin-Golomb, A. (2010). Gender differences in Parkinson’s disease: clinical characteristics and cognition. Mov Disord, 25(16), 2695–2703. doi:10.1002/mds.23388

Moisan, F., Kab, S., Mohamed, F., Canonico, M., Le Guern, M., Quintin, C., … Elbaz, A. (2016). Parkinson disease male-to-female ratios increase with age: French nationwide study and meta-analysis. J Neurol Neurosurg Psychiatry, 87(9), 952–957. doi:10.1136/jnnp-2015-312283

Morris, R. G., Schenk, F., Tweedie, F., & Jarrard, L. E. (1990). Ibotenate Lesions of Hippocampus and/or Subiculum: Dissociating Components of Allocentric Spatial Learning. Eur J Neurosci, 2(12), 1016–1028. doi:10.1111/j.1460-9568.1990.tb00014.x

Olsen, R. H., Johnson, L. A., Zuloaga, D. G., Limoli, C. L., & Raber, J. (2013). Enhanced hippocampus-dependent memory and reduced anxiety in mice over-expressing human catalase in mitochondria. J Neurochem, 125(2), 303–313. doi:10.1111/jnc.12187

Osterberg, V. R., Spinelli, K. J., Weston, L. J., Luk, K. C., Woltjer, R. L., & Unni, V. K. (2015). Progressive aggregation of alpha-synuclein and selective degeneration of lewy inclusion-bearing neurons in a mouse model of parkinsonism. Cell Rep, 10(8), 1252–1260. doi:10.1016/j.celrep.2015.01.060

Ostrerova, N., Petrucelli, L., Farrer, M., Mehta, N., Choi, P., Hardy, J., & Wolozin, B. (1999). alpha-Synuclein shares physical and functional homology with 14-3-3 proteins. J Neurosci, 19(14), 5782–5791. Retrieved from https://www.ncbi.nlm.nih.gov/pubmed/10407019

Pandey, S., & Srivanitchapoom, P. (2017). Levodopa-induced Dyskinesia: Clinical Features, Pathophysiology, and Medical Management. Ann Indian Acad Neurol, 20(3), 190–198. doi:10.4103/aian.AIAN_239_17

Pasanen, P., Myllykangas, L., Siitonen, M., Raunio, A., Kaakkola, S., Lyytinen, J., … Paetau, A. (2014). Novel alpha-synuclein mutation A53E associated with atypical multiple system atrophy and Parkinson’s disease-type pathology. Neurobiol Aging, 35(9), 2180 e2181-2185. doi:10.1016/j.neurobiolaging.2014.03.024

Polymeropoulos, M. H., Lavedan, C., Leroy, E., Ide, S. E., Dehejia, A., Dutra, A., … Nussbaum, R. L. (1997). Mutation in the alpha-synuclein gene identified in families with Parkinson’s disease. Science, 276(5321), 2045–2047. doi:10.1126/science.276.5321.2045

Proukakis, C., Dudzik, C. G., Brier, T., MacKay, D. S., Cooper, J. M., Millhauser, G. L., … Schapira, A. H. (2013). A novel alpha-synuclein missense mutation in Parkinson disease. Neurology, 80(11), 1062–1064. doi:10.1212/WNL.0b013e31828727ba

Prusiner, S. B. (2012). Cell biology. A unifying role for prions in neurodegenerative diseases. Science, 336(6088), 1511–1513. doi:10.1126/science.1222951

Rodriguez-Araujo, G., Nakagami, H., Takami, Y., Katsuya, T., Akasaka, H., Saitoh, S., … Kaneda, Y. (2015). Low alpha-synuclein levels in the blood are associated with insulin resistance. Sci Rep, 5, 12081. doi:10.1038/srep12081

Sasaki, A., Arawaka, S., Sato, H., & Kato, T. (2015). Sensitive western blotting for detection of endogenous Ser129-phosphorylated alpha-synuclein in intracellular and extracellular spaces. Sci Rep, 5, 14211. doi:10.1038/srep14211

Schaser, A. J., Osterberg, V. R., Dent, S. E., Stackhouse, T. L., Wakeham, C. M., Boutros, S. W., … Unni, V. K. (2019). Alpha-synuclein is a DNA binding protein that modulates DNA repair with implications for Lewy body disorders. Sci Rep, 9(1), 10919. doi:10.1038/s41598-019-47227-z

Schoch, K. M., & Miller, T. M. (2017). Antisense Oligonucleotides: Translation from Mouse Models to Human Neurodegenerative Diseases. Neuron, 94(6), 1056–1070. doi:10.1016/j.neuron.2017.04.010

Spillantini, M. G., Crowther, R. A., Jakes, R., Cairns, N. J., Lantos, P. L., & Goedert, M. (1998a). Filamentous alpha-synuclein inclusions link multiple system atrophy with Parkinson’s disease and dementia with Lewy bodies. Neurosci Lett, 251(3), 205–208. Retrieved from https://www.ncbi.nlm.nih.gov/pubmed/9726379

Spillantini, M. G., Crowther, R. A., Jakes, R., Hasegawa, M., & Goedert, M. (1998b). alpha-Synuclein in filamentous inclusions of Lewy bodies from Parkinson’s disease and dementia with lewy bodies. Proc Natl Acad Sci U S A, 95(11), 6469–6473. doi:10.1073/pnas.95.11.6469

Torres, E. R. S., Akinyeke, T., Stagaman, K., Duvoisin, R. M., Meshul, C. K., Sharpton, T. J., & Raber, J. (2018). Effects of Sub-Chronic MPTP Exposure on Behavioral and Cognitive Performance and the Microbiome of Wild-Type and mGlu8 Knockout Female and Male Mice. Front Behav Neurosci, 12, 140. doi:10.3389/fnbeh.2018.00140

Tran, H. T., Chung, C. H., Iba, M., Zhang, B., Trojanowski, J. Q., Luk, K. C., & Lee, V. M. (2014). Alpha-synuclein immunotherapy blocks uptake and templated propagation of misfolded alpha-synuclein and neurodegeneration. Cell Rep, 7(6), 2054–2065. doi:10.1016/j.celrep.2014.05.033

Ueda, K., Fukushima, H., Masliah, E., Xia, Y., Iwai, A., Yoshimoto, M., … Saitoh, T. (1993). Molecular cloning of cDNA encoding an unrecognized component of amyloid in Alzheimer disease. Proc Natl Acad Sci U S A, 90(23), 11282–11286. doi:10.1073/pnas.90.23.11282

van Putten, M. (2016). The use of hanging wire tests to monitor muscle strength and condition over time. Retrieved from http://www.treat-nmd.eu/downloads/file/sops/dmd/MDX/DMD_M.2.1.004.pdf

Vivacqua, G., Casini, A., Vaccaro, R., Fornai, F., Yu, S., & D’Este, L. (2011). Different sub-cellular localization of alpha-synuclein in the C57BL\6J mouse’s central nervous system by two novel monoclonal antibodies. J Chem Neuroanat, 41(2), 97–110. doi:10.1016/j.jchemneu.2010.12.003

Volpicelli-Daley, L. A., Luk, K. C., Patel, T. P., Tanik, S. A., Riddle, D. M., Stieber, A., … Lee, V. M. (2011). Exogenous alpha-synuclein fibrils induce Lewy body pathology leading to synaptic dysfunction and neuron death. Neuron, 72(1), 57–71. doi:10.1016/j.neuron.2011.08.033

Wang, C., Zhao, C., Li, D., Tian, Z., Lai, Y., Diao, J., & Liu, C. (2016). Versatile Structures of alpha-Synuclein. Front Mol Neurosci, 9, 48. doi:10.3389/fnmol.2016.00048

Weintraub, D., Siderowf, A. D., Potenza, M. N., Goveas, J., Morales, K. H., Duda, J. E., … Stern, M. B. (2006). Association of dopamine agonist use with impulse control disorders in Parkinson disease. Arch Neurol, 63(7), 969–973. doi:10.1001/archneur.63.7.969

Weitzner, D. S., Engler-Chiurazzi, E. B., Kotilinek, L. A., Ashe, K. H., & Reed, M. N. (2015). Morris Water Maze Test: Optimization for Mouse Strain and Testing Environment. J Vis Exp(100), e52706. doi:10.3791/52706

Wittmann, B. C., Schott, B. H., Guderian, S., Frey, J. U., Heinze, H. J., & Duzel, E. (2005). Reward-related FMRI activation of dopaminergic midbrain is associated with enhanced hippocampus-dependent long-term memory formation. Neuron, 45(3), 459–467. doi:10.1016/j.neuron.2005.01.010

Wu, L. G., Hamid, E., Shin, W., & Chiang, H. C. (2014). Exocytosis and endocytosis: modes, functions, and coupling mechanisms. Annu Rev Physiol, 76, 301–331. doi:10.1146/annurev-physiol-021113-170305

Xu, J., Wu, X. S., Sheng, J., Zhang, Z., Yue, H. Y., Sun, L., … Wu, L. G. (2016). alpha-Synuclein Mutation Inhibits Endocytosis at Mammalian Central Nerve Terminals. J Neurosci, 36(16), 4408–4414. doi:10.1523/JNEUROSCI.3627-15.2016

Zarranz, J. J., Alegre, J., Gomez-Esteban, J. C., Lezcano, E., Ros, R., Ampuero, I., … de Yebenes, J. G. (2004). The new mutation, E46K, of alpha-synuclein causes Parkinson and Lewy body dementia. Ann Neurol, 55(2), 164–173. doi:10.1002/ana.10795

Zhao, H. T., John, N., Delic, V., Ikeda-Lee, K., Kim, A., Weihofen, A., … Volpicelli-Daley, L. A. (2017). LRRK2 Antisense Oligonucleotides Ameliorate alpha-Synuclein Inclusion Formation in a Parkinson’s Disease Mouse Model. Mol Ther Nucleic Acids, 8, 508–519. doi:10.1016/j.omtn.2017.08.002

Zharikov, A., Bai, Q., De Miranda, B. R., Van Laar, A., Greenamyre, J. T., & Burton, E. A. (2019). Long-term RNAi knockdown of alpha-synuclein in the adult rat substantia nigra without neurodegeneration. Neurobiol Dis, 125, 146–153. doi:10.1016/j.nbd.2019.01.004

